# Inter-phylum circulation of a beta-lactamase - encoding gene: a rare but observable event

**DOI:** 10.1101/2023.11.10.566511

**Authors:** Rémi Gschwind, Marie Petitjean, Claudine Fournier, Julie Lao, Olivier Clermont, Patrice Nordmann, Alexander Mellmann, Erick Denamur, Laurent Poirel, Etienne Ruppé

**Author notes:** Corresponding author Etienne Ruppé, PharmD, PhD, Laboratoire de Bactériologie, Hôpital Bichat-Claude Bernard 46 rue Henri Huchard, 75018 Paris, France, E mail. Both authors contributed equally (Rémi Gschwind, Marie Petitjean).

## Abstract

Beta-lactam degradation by beta-lactamases is the most common mechanism of beta-lactam resistance in Gram-negative bacteria. Beta-lactamase encoding genes can be transferred between closely-related bacteria, but spontaneous inter-phylum transfers (between distantly related bacteria) has never been reported. Here, we describe an extended-spectrum beta-lactamase (ESBL)-encoding gene (*bla*_MUN-1_) shared between the Peudomonadota and Bacteroidota phyla.

An *Escherichia coli* strain was isolated from a patient in Münster (Germany). Its genome was sequenced (Illumina and Nanopore). The ESBL encoding gene was cloned and the corresponding enzyme was characterised. Distribution of the gene among bacteria was studied with BLASTN using RefSeq Genomes databases. Frequency of its closest homolog in the Global Microbial Gene Catalog (GMGC) was also analysed.

The *bla*_MUN-1_ gene found in the *E. coli* strain, encoded for an Ambler subclass A2 beta-lactamase with 82.2% amino acid identity to TLA-1 and it was found to confer an ESBL phenotype. *bla*_MUN-1_ was found in four copies, two chromosomal copies and two located on a phage-plasmid p0111. Each copy was found on a 7.6kb genomic island associated with mobility. *bla*_MUN-1_ was found distributed among the Bacteroidales order and in *Sutterella wardsworthensis* (Pseudomonadota). Its closest homolog in the GMGC was found predominantly and frequently in the Human gut sub-catalog (found in 26.8% of the samples).

This is the first reported case of inter-phylum transfer of an ESBL-encoding gene, between the Bacteroidota and Pseudomonadota phyla. While the gene was frequently found in the human gut, inter-phylum transfer was rare, suggesting that inter-phylum barriers are strong but not impassable.

## Introduction

Beta-lactamases refer to enzymes catalyzing the hydrolysis of the beta-lactam ring, thereby inactivating the antibiotic properties of the molecule. According to the classification proposed by Ambler (1), beta-lactamases are distributed into four classes, with class A, C, and D beta-lactamases using a serine residue to catalyze the reaction, while class B beta-lactamases (also referred to as metallo-beta-lactamases) use one or two atoms of ionized zinc to do so (2). Beta-lactamases are widely spread among bacteria irrespective of their pathogenic or commensal, environmental, animal or human traits. They are the main resistance mechanism by which several pathogens resist to beta-lactams, especially for Gram-negative bacilli from the Pseudomonadota phylum such as Enterobacterales, *Pseudomonas aeruginosa* and *Acinetobacter baumannii*.

While some Enterobacterales intrinsically harbor beta-lactamases, the biggest threat to health is due to the acquisition and exchange by pathogens of beta-lactamases–encoding genes, especially those encoding for extended-spectrum beta-lactamases (ESBL) and carbapenemases, which inactivate most available beta-lactam antibiotics. Enterobacterales have indeed the capacity to share mobile genetic elements (MGE) embedding antibiotic resistance genes (ARGs) via conjugation, however how the first move from the original host of the ARG and Enterobacterales species is not known in most instances. In a recent work, Ebmeyer *et al.* described the origin of 37 ARGs found in Enterobacterales and provide evidence for the original gene-providing species for 27 groups of ARGs (3). Strikingly, 36/37 of transfer events occurred among Pseudomonadota (to which Enterobacterales belong), with *tet(X)* being a notable exception as no conclusive evidence could be found. Indeed, it was proposed to originate from *Sphingomonas*, a genus from the Bacteroidota phylum (4). However, further analysis revealed that the gene was only present in 4 out of 26 *Sphingomonas spp.* genomes and it was always associated with mobility (mobilizable transposon-like element), which weakens the assumption that *Sphingomonas* is the donor genus origin of *tet(X)*. Yet, the identification of *tet(X)* in bacteria from two distinct phyla showed that such transfer could occur.

This part of shadow regarding ARG-providing species to Enterobacterales pointed at the intestinal microbiota as a potential reservoir of ARGs (5). The dominant fraction of the intestinal microbiota is made of strict anaerobic bacteria, the most abundant being from the phyla Firmicutes and Bacteroidota. They indeed possess a vast diversity of ARGs including some encoding beta-lactamases (6), many of which proven to be functional when transferred to *Escherichia coli* (7). However, ARGs from commensal anaerobic bacteria strongly differ from those found in Enterobacterales, stressing that if ARG transfers from anaerobic bacteria to Enterobacterales actually occurred, they would be particularly rare or would not persist so it would go unseen from the scientific community (8). Hence, inter-phylum transfers of ARGs seem to be exceptional to observe outside laboratory conditions.

In a recent work, we searched for ARGs in 70k *E. coli* genomes from the Enterobase using ARG databases including ARGs from intestinal strict anaerobic bacteria (9, 10). We could identify four ARGs presumably originating from non-Pseudomonadota, including a beta-lactamase encoding gene also found in bacteria from the Bacteroidota phylum and that we propose to characterize in the present work.

## Results

### Phenotypic characterization

Analyzing 70,301 *E. coli* genomes available from the Enterobase, one was identified as possessing a beta-lactamase - encoding gene which was only found in the ResFinderFG and Mustard databases (6, 10, 11). The strain of interest was an *E. coli* strain belonging to the A phylogroup, sequence type (ST) 744/2 according to the Warwick University/Pasteur Institut schemes, respectively, serotype Onovel132:H10, *fimH* allele 54. It was isolated in 2015 from a wound infection in a patient hospitalized at the University Hospital of Münster, Germany (12). Of note, subcultures in LB medium yielded two distinct morphotypes. The first type corresponded to white and regular shaped colonies while the second was represented by greyish and less regular colonies (Figure 1). Both morphotypes were maintained in subsequent cultures.

**Figure 1:**
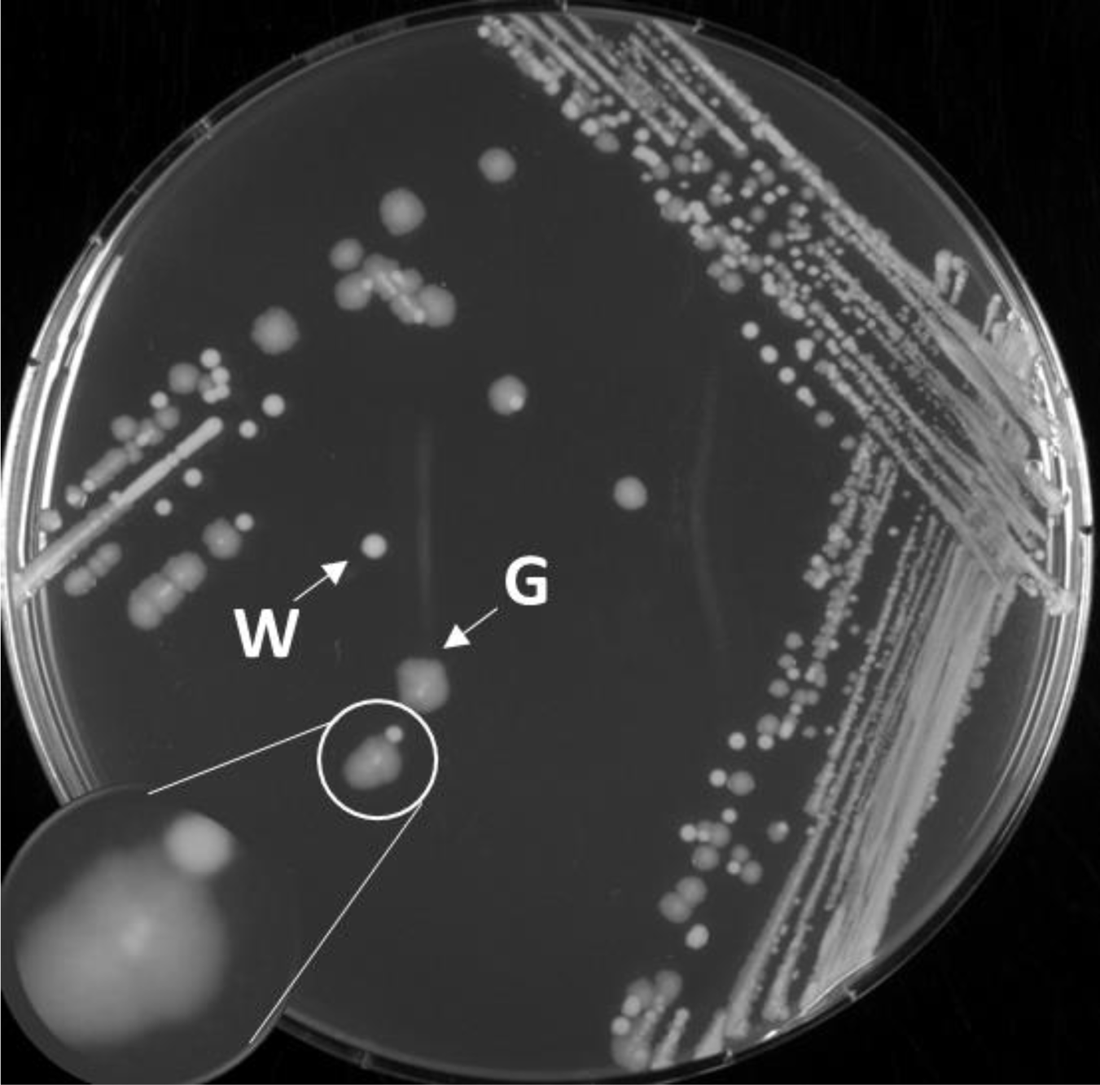
Morphological aspects of the two types of colonies (W: white colonies, G: grey colonies) observed after streaking the strain on LB media.

From the antibiotic susceptibility testing performed by the disk diffusion method, the *E. coli* strain characterized by white and regular colonies displayed an ESBL phenotype with synergies being observed between clavulanic acid, cefotaxime, cefepime, and aztreonam (Supplementary Figure 1). Particularly, the strain showed a high level of resistance to cefuroxime, ceftazidime and aztreonam with MIC > 256 µg/mL (Table 1), and more surprisingly, to temocillin (MIC > 256 µg/L). It remained susceptible to carbapenems and cefoxitin, and to beta-lactams-beta-lactam inhibitor combinations (clavulanic acid, tazobactam and avibactam). Besides, the strain was resistant to cotrimoxazole and fluoroquinolones but was susceptible to aminoglycosides (gentamicin, netilmicin, amikacin and tobramycin). The same phenotype was observed for grey colonies, except for some beta-lactam antibiotics (aztreonam, cefuroxime, cefotaxime, ceftazidime and piperacillin) against which grey colonies were slightly less resistant (as observed from inhibition diameters).

**Table 1:** Minimal inhibitory concentrations of the white colonies, the *E. coli* TOP10 cloned or not cloned with the *bla*_MUN-1_ gene and the kinetic parameters of purified MUN-1 beta-lactamase. ND: not done.

| Beta-lactam | MIC (µg/mL) for *Escherichia coli* |  |  | Kinetic measurement |  |  |  |
| --- | --- | --- | --- | --- | --- | --- | --- |
| | Clinical strain | TOP10 (pBLA-x) | TOP10 | $K_{cat}$ (s<sup>-1</sup>) | $K_m$ (µM) | $K_{cat}/K_m$ (µM<sup>-1</sup>s<sup>-1</sup>) | $K_i$ (µM) |
| Amoxicillin | > 256 | > 256 | 4 | ND | ND | ND | ND |
| Amoxicillin + clavulanic acid | 4 | 4 | 4 | - | - | - | - |
| Ampicillin | ND | ND | ND | 290 | 280 | 1 | ND |
| Piperacillin | > 256 | > 256 | 2 | < 0.01 | ND | ND | 0.0072 |
| Piperacillin + tazobactam | 4 | 4 | 2 | - | - | - | - |
| Penicillin G | ND | ND | ND | 210 | 95 | 2.2 | ND |
| Temocillin | 256 | 256 | 4 | ND | ND | ND | ND |
| Ticarcillin | ND | ND | ND | < 0.01 | ND | ND | 0.0036 |
| Cephalothin | ND | ND | ND | 30 | 15 | 2 | ND |
| Cefuroxime | > 256 | > 256 | 0.5 | ND | ND | ND | ND |
| Ceftriaxone | 16 | 16 | 0.12 | ND | ND | ND | ND |
| Cefotaxime | 4 | 2 | 0.12 | < 0.01 | ND | ND | ND |
| Ceftazidime | > 256 | > 256 | 0.25 | < 0.01 | ND | ND | 0.58 |
| Ceftazidime + avibactam | 0.06 | 0.03 | 0.25 | - | - | - | - |
| Cefepime | 4 | 2 | 0.06 | < 0.01 | ND | ND | ND |
| Ceftolozane + tazobactam | 0.06 | 0.06 | 0.06 | - | - | - | - |
| Aztreonam | > 256 | > 256 | 0.03 | 2 | 45 | 0.04 | ND |
| Imipenem | 0.25 | 0.25 | 0.25 | < 0.01 | ND | ND | 0.0013 |
| Meropenem | 0.03 | 0.03 | 0.03 | < 0.01 | ND | ND | 0.019 |
| Ertapenem | 0.03 | 0.03 | 0.03 | < 0.01 | ND | ND | 0.017 |

### Beta-lactamase characterization

The beta-lactamase amino-acid sequence was studied and compared to other amino acid sequences of other beta-lactamases (Figure 2). It was shown to be an Ambler subclass A2 beta-lactamase (13). Looking at the cophenetic distance, the closest beta-lactamase to MUN-1 were TLA-1 (1.09; 82.2% amino acid identity), CepA (1.15; 72.3% amino acid identity) and VEB-1 (1.34; 71.2% amino acid identity).

**Figure 2:**
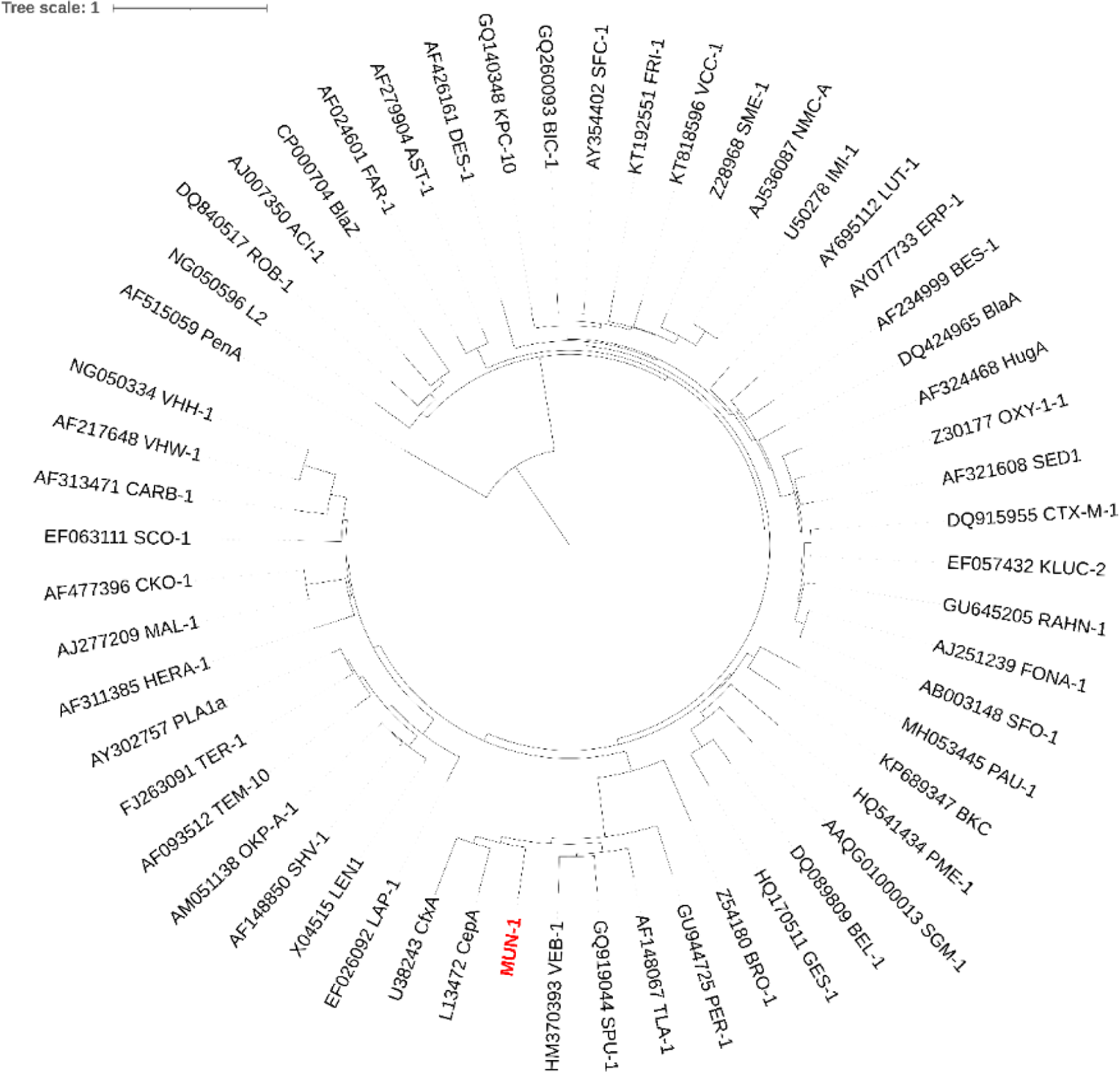
Phylogenetic tree of representative beta-lactamases found in the bacterial realm including the MUN-1 beta-lactamase (in red). Phylogenetic tree was rooted on PenA (found in the genus *Burkholderia*) which is distantly related from all the other beta-lactamases.

The *bla*_MUN-1_ gene was cloned and expressed in *E. coli* TOP10. A similar resistance phenotype to the original strain was observed (Table 1). No hydrolysis could be detected with ticarcillin, piperacillin and ceftazidime as substrates, although increased MIC values were observed for the recombinant strain (Table 1). Conversely, a significant hydrolysis was observed with aztreonam in consistency with the high-level resistance observed in the clinical strain and the transformant. As for the susceptibility to beta-lactamase inhibitors, the most potent one was clavulanic acid (IC50 0.32 nM), followed by tazobactam (IC50 0.8 nM) and avibactam (IC50 3.8 nM).

### Molecular characterization

White and grey *E. coli* strain colonies were sequenced using short read and long read technologies to determine the *bla*_MUN-1_ gene locations. The hybrid assembly yielded two contings for the grey colonies and 3 contigs for the white colonies (Supplementary Table 1).

A circular bacterial chromosome of 4,764,212 and 4,762,657 bp was identified for the grey and white colonies respectively. The ARG and virulence gene content was similar in both morphotypes (Supplementary Table 2 and 3). Of note, *dfrA17*, *sul1*, *sul2*, *qacE*, *mph(A)*, *bla*_TEM-1_, *aph(3’)-Ia*, *aph*(*6*)*-Id*, *aph(3’’)-Ib*, *aadA5* and *catA1*) were found on an antibiotic resistance genomic island consisting in an inserted plasmid sequence IncQ1 surrounded by 10 insertion sequences (Supplementary Figure 2). Additionally, a tetracycline resistance gene *tet(B)* was also identified but outside this antibiotic resistance genomic island. The only difference in the bacterial chromosome between the two morphotypes consisted in the presence of two additional insert sequences (0.78 kb each containing IS1 family transposase and IS1 family IS1SD transposase ORF B) in the chromosome from the grey strain. First insertion was responsible for the shortening of a *mshA* gene encoding for a glycosyltransferase. The second one was inserted inside a L,D-transpeptidase encoding gene. Regarding the *bla*_MUN-1_ gene identified in the previous study, it was found in two copies on the bacterial chromosome outside the antibiotic resistance genomic island. Each *bla*_MUN-1_ copy was found on a 7.6 kb genomic island (Figure 3) annotated with 5 additional ORFs encoding for site-specific integrase, a helix turn helix crp-type domain-containing protein, a helicase, a DNA primase and a plasmid recombination enzyme. The first 7.6 kb genomic island containing *bla*_MUN-1_ gene was located at 59.85 min on the *E. coli* genetic map. It was surrounded by genes involved in the GABA degradation pathway allowing GABA use as a nitrogen source (*gabD*, *gabT*, *gabP*) and genes involved in rescue of stalled ribosomes mediated by trans-translation (*ssrA*, *smpB*). The second 7.6 kb genomic island containing the *bla*_MUN-1_ gene was located at 93.42 min, between the *adiA* gene (encoding a biodegradative arginine decarboxylase) and the *melAB* operon (involved in melibiose metabolism). The only shared characteristic found at each border of each 7.6 kb genomic island was their low GC content (mean of 34.3% GC in the 200 bp flanking each 7.6 kb genomic island copy while 50.6% GC was observed for the entire chromosome).

**Figure 3:**
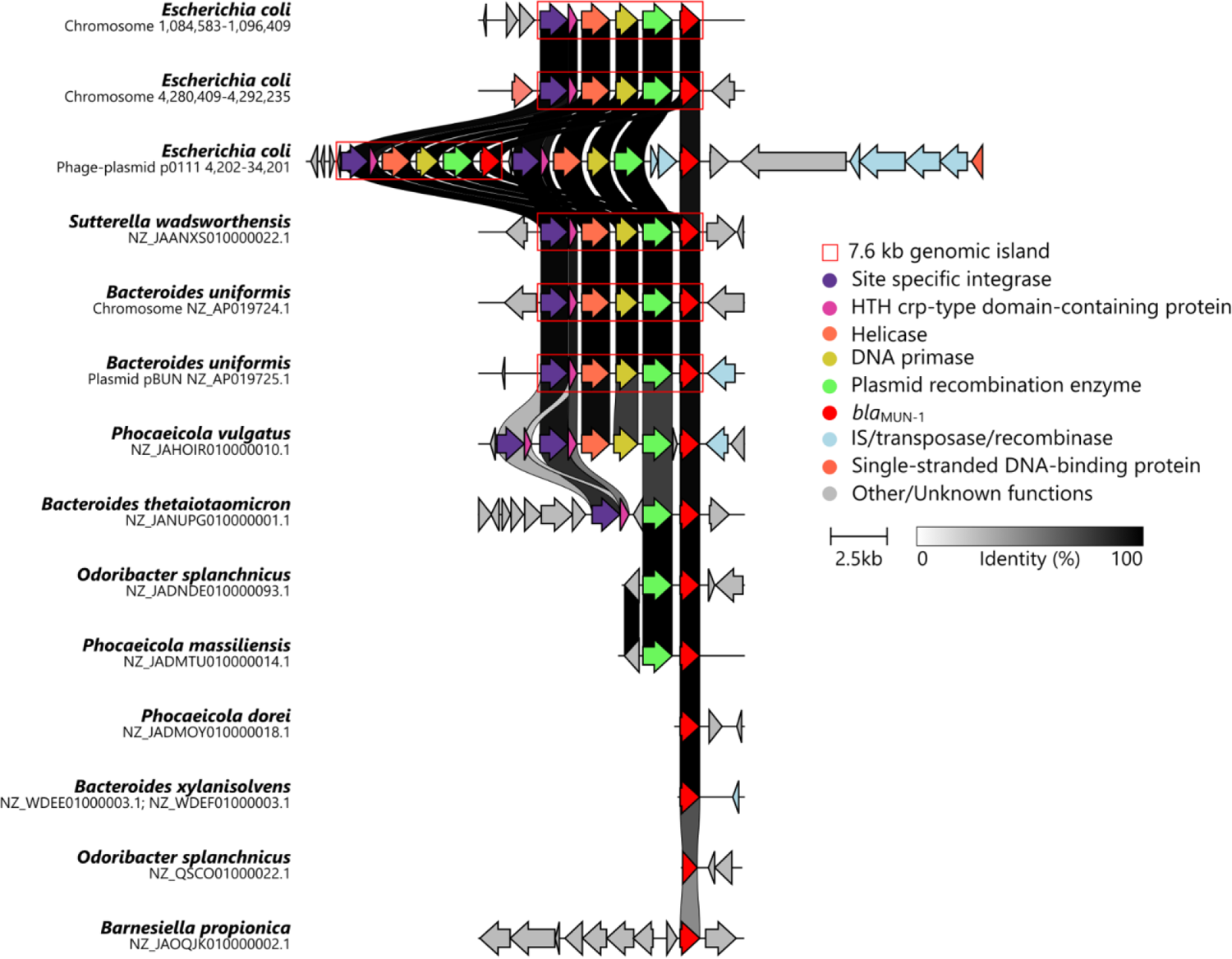
Genetic contexts showing the environment of the *bla*_MUN-1_ gene in different species. The three first lines describe the genetic contexts of each copy of the *bla*_MUN-1_ gene in the *E. coli* strain. Next, illustrative representatives from the RefSeq Genomes database were chosen for the following reasons: *S. wadsworthensis* was the only other Pseudomonadota found to bear the *bla*_MUN-1_ gene; *B. uniformis* was the only genome in which copies of *bla*_MUN-1_ gene were found on a chromosome and on a plasmid; *P. vulgatus*, *B. thetaiotaomicron*, *O. splanchnicus*, *P. massiliensis*, *P. dorei*, *B. xylanisolvens* and *B. propionica* were chosen as they were the only genomes showing a genetic contexts that seemed to be different from the 7.6 kb genomic island. The red box delineates the 7.6 kb genomic island described in this work. Colors in arrows correspond to the function of each gene. A nucleic acid identity percentage between adjacent lines is displayed with a grey scale.

Then, a 127 kb circular p0111 phage-plasmid was found only in the white colonies (Supplementary Table 1). Two copies of the *bla*_MUN-1_ gene were identified on the p0111 phage-plasmid. One copy was found on a 7.6 kb genomic island with 100% nucleic acid identity with the ones found on the chromosome. Adjacent to this genomic island, a second 7.6 kb genomic island carrying the other gene copy was found. However, this island was different due to the insertion of two IS (IS3 family transposase ISEc52) between the *bla*_MUN-1_ gene and the plasmid recombination enzyme encoding gene. As in the bacterial chromosome, low GC content was found at the border of each 7.6 kb genomic island (Mean of 31.5% while 46.6% on the whole phage-plasmid). Next to the two 7.6 kb genomic islands, a 11 kb DNA fragment (containing a FRG domain-containing protein encoding gene, a AAA domain-containing protein encoding gene, two integrase encoding genes, a recombinase encoding gene and a transposase encoding gene) was found to be shared between the p0111 and the chromosome suggesting recombinations between the p0111 and the chromosome. Besides the bacterial chromosome, a circular IncFII plasmid of 60 kb was found for each morphotype. Of note, it did not embed any ARG.

### Distribution of the bla_MUN-1_ gene

We searched for *bla*_MUN-1_ using RefSeq Genomes databases from NCBI (>70% nucleic acid identity, 80% coverage) and BLASTN (14). A total of 125 hits were obtained, with the *bla*_MUN-1_ gene being found in 28 species with 100% nucleic acid identity and coverage (Supplementary Table 4). Most of the species were from the Bacteroidota phylum and more precisely from the Bacteroidales order. A unique hit was obtained in the Pseudomonadota phylum with *Sutterella wadsworthensis*. Among all the hits, we could not determine whether the sequence holding the *bla*_MUN-1_ gene was chromosomal or plasmidic, one exception being a *Bacteroides uniformis* genome (AP019724.1 and AP019725.1) bearing two copies of the 7.6 kb genomic island (containing the the *bla*_MUN-1_ gene), one being on a plasmid surrounded by sequences annotated as IS256 family transposase and site-specific integrase. Of note, two sequences on this plasmid were annotated as phage protein. Few species displayed variation in the sequence of the *bla*_MUN-1_ gene. Hits without 100% identity were obtained in *Bacteroides salyersiae*, *Bacteroides xylanisolvens*, *Parabacteroides distasonis*, *P. stercorea* and *Phocaeicola vulgatus* (97.0-99.9% nucleic acid identity). Additionally, *Barnesiella propionica* was shown to bear a gene with 71.9% nucleic acid identity and 96% coverage to *bla*_MUN-1_ gene. As in our *E. coli* strain of interest, the gene was occasionally found in several copies (maximum of 6 copies/genome in *B. uniformis*). Of note, *bla*_MUN-1_ was not found in all genomes available from a given species in any species (Supplementary Table 4). Additionally, MGnify and the global microbial gene catalog (GMGC) databases were used to analyze the distribution of the *bla*_MUN-1_ gene (15, 16). The same results were found, with most of the hits in bacteria from the Bacteroidales order (Supplementary Table 5 and 6). We analysed the distribution of the best match obtained with GMGC (GMGC10.047_051_980.UNKNOWN - *Prevotellamassilia timonensis* - 86.8% cover and 100% identity). It was found in several sub-catalogs (Figure 4). However, it was mainly found in the Human gut sub-catalog where it was found in 26.8% of the samples, with a mean relative abundance of 104.5/10M reads (median: 12, min: 0, max: 5371).

**Figure 4:**
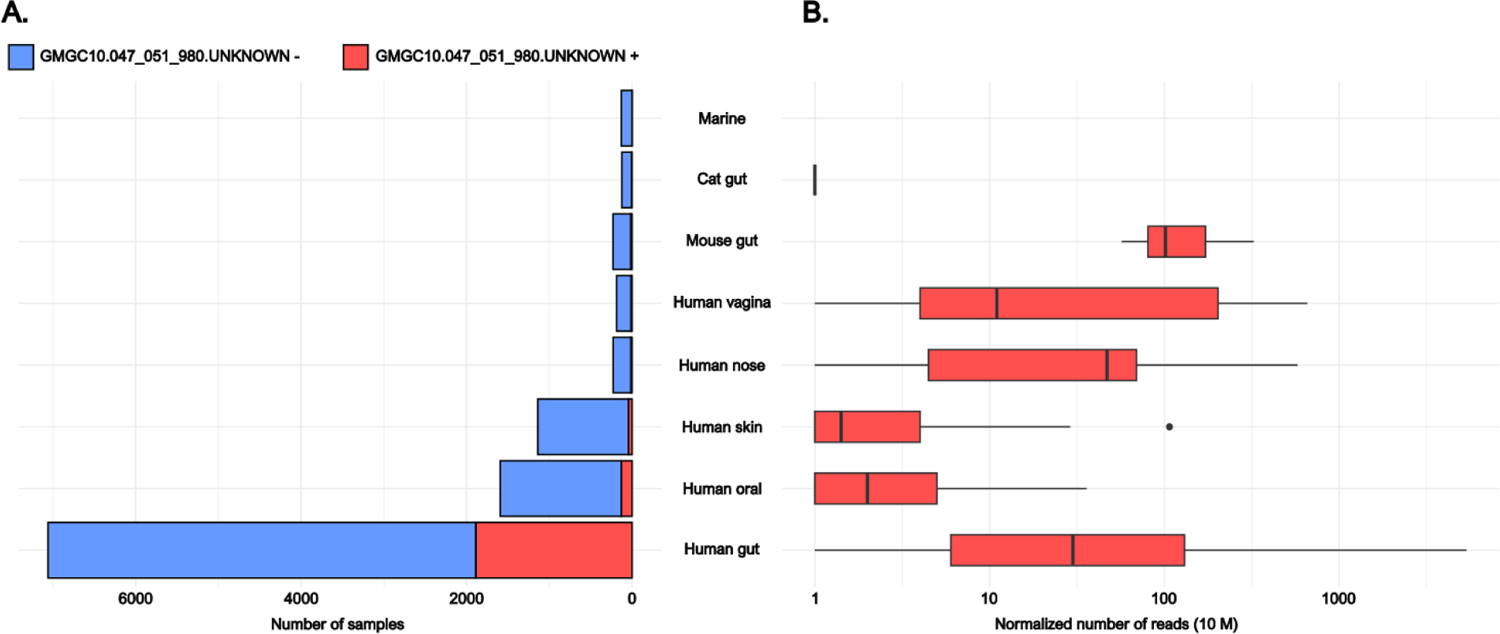
Frequency of the GMGC unigene GMGC10.047_051_980.UNKNOWN (100% identity and 86.8% cover in amino acid with the *bla_MUN-1_* gene) in unigene sub-catalogs where it is found and the associated number of mapped reads. A: Number of samples found in each sub-catalog containing (in red) or not (in blue) the GMGC unigene GMGC10.047_051_980.UNKNOWN. B: Normalized number of reads (out of 10 million reads) mapping onto the GMGC unigene GMGC10.047_051_980.UNKNOWN in each GMGC sub-catalog where it was found. The normalization takes into account the size of the gene and the number of reads in each sample from the sub-catalogs.

The *bla*_MUN-1_ gene environment was analyzed in every hit using Bakta annotation. It was always found to be held by the 7.6 kb genomic island which was also found in the *E. coli* of interest (Figure 3) except for 9 out of 125 hits which were characterized by specific genetic contexts. Two hits were characterized by different 7.6 kb genomic islands. In a *P. vulgatus* strain (NZ_JAHOIR010000010.1), the *bla*_MUN-1_ gene was found on a 7.6 kb genomic island with 82.0% nucleic acid identity. Then, in a *Bacteroides thetaiotaomicron* strain (NZ_JANUPG010000001.1), it was found on a shorter version of the 7.6 kb genomic island (63% cover) which consisted of the *bla*_MUN-1_ gene (100% cover, 100% identity), the plasmid recombination enzyme (100% cover, 99.9% identity) and the site-specific integrase encoding genes (99% cover, 80.60% identity). Then, all other genetic contexts were specific as they were found at the border of a contig, meaning it was not possible to identify the 7.6 kb genomic island (NZ_WDEF01000003.1, NZ_WDEE01000003.1, *B. xylanisolvens*; NZ_JADNDE010000093.1, NZ_QSCO01000022.1 *Odoribacter splanchnicus*; NZ_JADMTU010000014.1 *Phocaiecola masssiliensis*; NZ_JADMOY010000018.1 *Phocaeicola dorei*; NZ_QRNO01000005.1 *P. stercorea*). Finally, one last hit was found in *B. propionica* which was the only hit not associated with 7.6 kb genomic island but also with a lower nucleic acid identity regarding the *bla*_MUN-1_ gene sequence (71.9%).

To show the phylogenetic distance between all the *bla*_MUN-1_ bearing species, we built a phylogenetic tree using the 16S rRNA encoding gene of each species (Figure 5). We observed that from *E. coli*, the closest species holding the *bla*_MUN-1_ gene were *S. wadsworthensis* (cophenetic distance 0.22) and both species from the *Alistipes* genus (cophenetic distance 0.40). The next closest species was the species we used to root the phylogenetic tree, *Gemmata massiliana* (Planctomycetota phylum; cophenetic distance 0.43) which do not bear the *bla*_MUN-1_ gene. The most distant species bearing *bla*_MUN-1_ was *Prevotella stercorea* (cophenetic distance 0.61).

**Figure 5:**
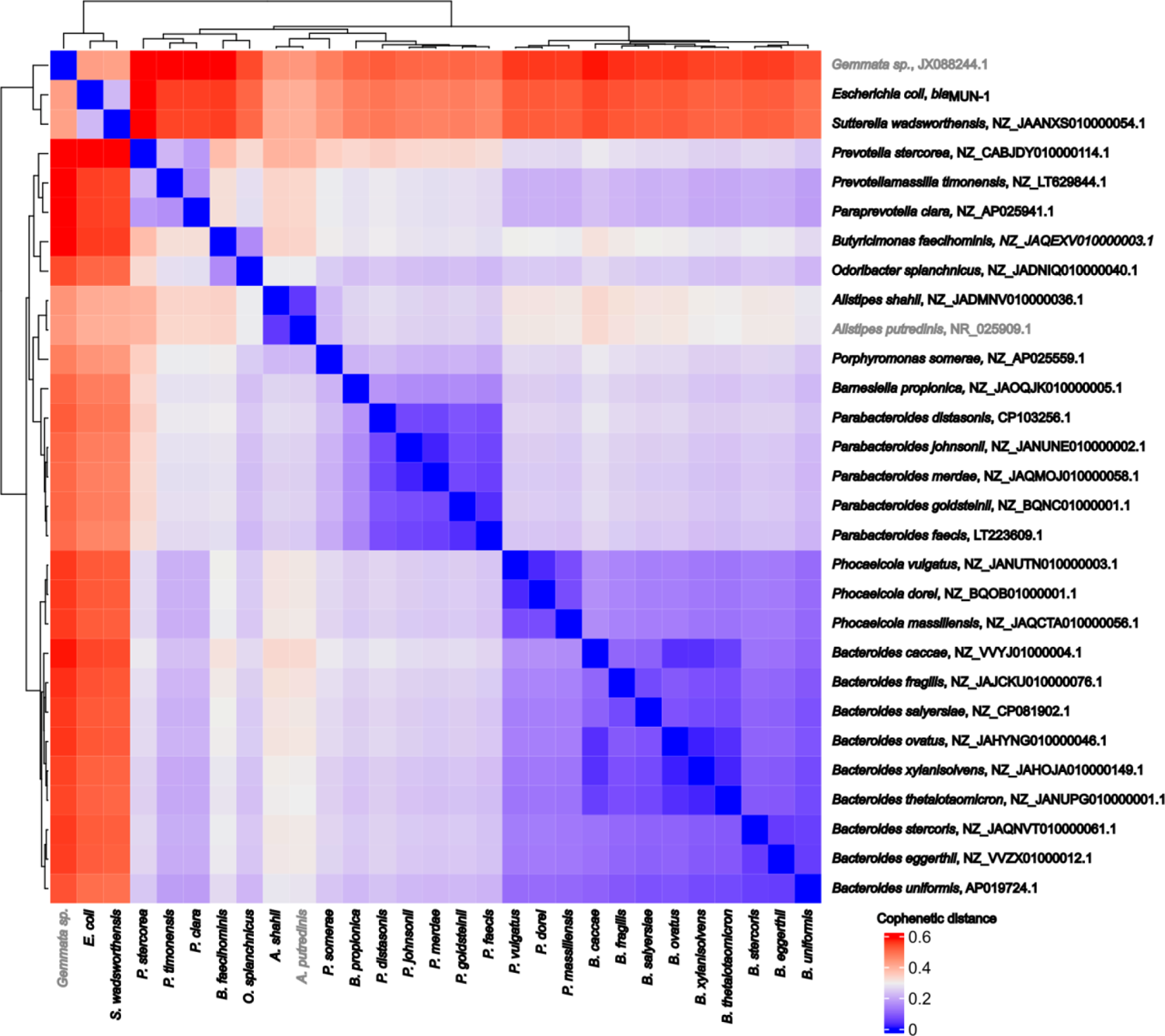
Cophenetic distance between species holding the *bla*_MUN-1_ gene based on the 16S rRNA encoding gene. Heatmap represents the cophenetic distance between species. If no strain holding the *bla*_MUN-1_ gene from the species was found to hold a 16S rRNA encoding gene, 16S rRNA encoding gene was retrieved from strains that do not hold the *bla*_MUN-1_ gene (This was the case for *A. putrenidis* in grey). *Gemmata sp.* did not hold a *bla*_MUN-1_ gene in its genome but its 16S rRNA encoding gene was used to root the phylogenetic tree.

## Discussion

A *bla*_MUN-1_ gene encoding for an ESBL was found to be shared between Bacteroidota phyla and Pseudomonadota phyla, suggesting that an inter-phylum transfer of ESBL encoding gene is possible.

The characterized MUN-1 beta-lactamase was an Ambler subclass A2 beta-lactamase with an ESBL phenotype (13). Notably, it conferred resistance to several beta-lactam antibiotics, including temocillin, which is unusual among class A beta-lactamases (17). The closest beta-lactamases were TLA-1 (82.2% amino acid identity) and CepA (72.3% amino acid identity). CepA is a characteristic beta-lactamase of *Bacteroides* genus and the TLA-1 encoding gene was initially described on a plasmid found in *E. coli* (18–20). Yet, the closest beta-lactamase encoding gene to TLA-1 encoding gene was the CME-1 beta-lactamase encoding gene from *Chryseobacterium meningosepticum*, also a member of the Bacteroidota phylum. However, TLA-1 and CME-1 encoding genes shared only 50% nucleic acid identity, making it difficult to determine the origin of TLA-1.

The distribution of *bla*_MUN-1_ among bacteria was studied and it was found mostly in bacteria from the Bacteroidales order and in one *S. wadsworthensis* genome (Pseudomonadota phylum). This suggests that inter-phylum transfer of the *bla*_MUN-1_ gene has occurred at least once. The genetic contexts around the gene were most often associated with the 7.6 kb genomic island that was conserved (100% nucleic acid identity), indicating that the inter-phylum transfer occurred relatively recently. Interestingly, all of the species identified using RefSeq Genomes in this study are either from sequencing projects linked to the intestinal microbiota or are characteristic species of the intestinal microbiota. In addition, by studying the distribution of the gene homologous to the *bla*_MUN-1_ gene in the GMGC catalogue, it was shown that it was not only present mainly in the Human gut sub-catalogue, but it was also found in more than a quarter of the Human gut samples. This gene is therefore frequent but the inter-phylum transfer seems to be rare suggesting that the phylum barrier for ARG transfer is efficient but can be overcome.

This paper has several limitations. As with TLA-1 encoding gene, origin of *bla*_MUN-1_ could not be determined because it was always found associated with mobility and in a small number of the species’ genomes (meaning that the prevalence of the gene in the species is low). Moreover, only hypotheses can be drawn about how the gene was transferred between phyla. We cannot state which Bacteroidales or Pseudomonadota bacteria were involved in this transfer. *A. putredinis* and *S. wadsworthensis* could be good candidates as they share the lowest cophenetic distance between Bacteroidota and Pseudomonadota 16S rRNA encoding genes. We also cannot state which MGE was involved. The *E. coli* strain was shown to possess two copies of the gene on a p0111 phage-plasmid that might be involved. Phage-plasmids are prophages having functions associated with plasmids and they were already shown to carry ARGs. p0111 is part of the P1 subgroup, a subgroup specific to *E. coli* which was not found in Bacteroidota phyla so far (21, 22). *bla*_MUN-1_ gene was also found on a plasmid in *B. uniformis* which could have participated in this inter-phylum transfer as well. It would have been possible to conduct experiments in a controlled environment to show the validity of these hypotheses. *In vitro* experiments showed that transformation of *E. coli* with a plasmid coming from *Bacteroides fragilis* was possible while conjugation between these two species was not (23). Yet, *in vitro* experiments between two strains do not necessarily reflect what happens *in vivo* in a complex bacterial ecosystem. Inter-phylum transfer of DNA, including conjugation between Bacteroidota and Pseudomonadota, was shown to be possible within complex bacterial communities (8, 23, 24). Thus, it is important to note that these *in vitro* experiments would not have proven how the *bla*_MUN-1_ gene was actually transferred between Bacteroidales and Pseudomonadota.

The *E. coli* strain of interest showed two morphotypes, one of which had an additional p0111 phage-plasmid carrying two additional copies of the *bla*_MUN-1_ gene. These repetitive regions and potential mixed strains add complexity to the analysis of sequencing data. Short and long read sequencing technology helps in read assembly but special care still needs to be taken in the description of all observed phenotypes. The biggest gap that needs to be filled concerns the general lack of knowledge regarding anaerobic bacteria. The data we are gradually collecting through sequencing and further analysis using computational methods will provide more insights into the true origin of the *bla*_MUN-1_ gene and help us better understand how this rare event occurred.

Here was the first to our knowledge, evidence of a shared ESBL-encoding gene between Bacteroidota and Pseudomonadota phyla. This observation shows that even though ARG transfers occur in closely-related species, a transfer between distantly-related species can spontaneously occur. How such transfer actually occurred and why it has not widely spread subsequently remains to be answered.

## Material and methods

### Bioinformatic analyses

In our previous work (9), ARGs were searched in 70,301 *E. coli* genomes available from the Enterobase^10^ using Diamond and the following ARG databases: AMRFinder (2019-04-29 version), Mustard (2017-09-30 version) and ResFinderFG (2016-11-01 version) (6, 11, 25, 26). A hit was considered as a potential ARG if the nucleic acid identity × coverage ≥ 0.64. All redundancies between the hits obtained in ≥2 databases were removed. The ARG families were picked according to the Mustard website (mgps.eu/Mustard). Among all hits, we identified a putative beta-lactamase encoding gene sharing 100% nucleic acid identity with a beta-lactamase-encoding gene from ResFinderFG (beta_lactamase|KU546399.1|feces|AMX, 100% identity, 93.9% cover) and Mustard (MC3.MG60.AS1.GP1.C4251.G1). We propose the name *bla*_MUN-1_ with regards to the city where the original strain was collected (Münster).

### Strain characterization

The *E. coli* strain with the beta-lactamase-encoding gene was re-tested for antibiotic susceptibility by the disk diffusion method on Mueller-Hinton agar according to the CASFM/EUCAST guidelines (2022 v1.0) and re-sequenced using Illumina (San Diego, CA) and Oxford Nanopore (Oxford Nanopore Technologies, UK) chemistries (Flongle R9.4.1) for obtaining short and long reads, respectively. The quality of Illumina and Nanopore reads were assessed using FastQC (v0.11.9). Trim galore (v0.6.7) was used to remove Illumina adapters and trimmed reads with a quality threshold of 30. The hybrid assembly of Illumina and Nanopore reads was performed using Unicycler (v0.4.9b) (27). The phylogroup of the strain was performed using the ClermonTyping (v23.06.05) tool and the sequence type with MLST (v2.19.0). Serotype and virulence genes were characterized using the ABRicate (1.0.0) software and the ecoh and a home-made database respectively. The *fimH* allele was determined using FimTyper (v1.1). ARGs were identified using the Diamond software (v2.1.8) and the ResFinder database (v4.0) (28). PlasmidFinder was used to characterize plasmid incompatibility groups. Contigs were annotated using Bakta (v1.8.2) (29, 30).

### Distribution of the bla_MUN-1_ gene

The distribution of *bla*_MUN-1_ and potential variants was assessed using BLASTN (>70% identity, 80% coverage) online with RefSeq Genomes database from NCBI (as of August 24^th^ 2023) (14). In each genome homologs of *bla*_MUN-1_ were found, its genetic environment was described using Bakta (v1.8.2) annotation and Clinker (31). One RefSeq genome containing the *bla*_MUN-1_ gene of each species was randomly chosen and a copy of its 16S rRNA encoding gene was retrieved. If no 16S rRNA encoding gene were available in any RefSeq genome from a species containing the *bla*_MUN-1_ gene, the 16S rRNA encoding gene was taken from another RefSeq genome of the species. The 16S rRNA encoding genes were used for alignment with MAFFT (v7.407) (32). To assess the phylogenetic distance of each species bearing the *bla*_MUN-1_ gene, a phylogenetic tree was made using IQ-TREE (v1.6.9, with ultrafast bootstrap and general time reversible model) (33, 34). Additionally, *bla*_MUN-1_ was also searched in the global microbial gene catalog (GMGC) and in MGnify (15, 16). The distribution and frequency of the best hit obtained with GMGC was also analyzed in the catalog.

### Comparison of MUN-1 to known beta-lactamases

The *bla*_MUN-1_ gene was translated into protein and aligned with other beta-lactamases retrieved from the ResFinder (v4.0) database using MAFFT (v7.407). To assess the phylogenetic distance between each beta-lactamases, another phylogenetic tree was made using IQ-TREE (v1.6.9, with ultrafast bootstrap and general time reversible model).

### Mating and cloning experiments

The *bla*_MUN-1_ gene was cloned into pTOPO-kanR vector using the pCR-Blunt TOPO cloning kit (Invitrogen) using specific primers (MUN Fw: 5’-GAG CAT CCG CTT CTT TGG GC-3’ and MUN Rv: 5’-ACG CAC ACT TTT CCG ATA TG-3’) spanning the full gene in order to express the whole protein. The resulting recombinant plasmid was transformed by heat shock into *E. coli* TOP10 (pTOPO/*bla*_MUN-1_).

### Protein and enzymatic characterization

Purification of the MUN-1 beta-lactamase was carried out by ion-exchange chromatography and its molecular mass was determined by SDS-12% PAGE (GeneScript) analysis. Purified beta-lactamase was used for kinetic measurements. IC50 values were determined for clavulanic acid, tazobactam, and avibactam (detailed protocol in the supplementary materials).

## Data summary

Illumina and nanopore reads were deposed under the BioProject PRJNA694822.

## Acknowledgements

We thank Prof. Alain Philippon, Prof. Eduardo Rocha and Dr Eugen Pfeifer for helpful discussions.

## Supplementary materials

**Supplementary Table 1:**
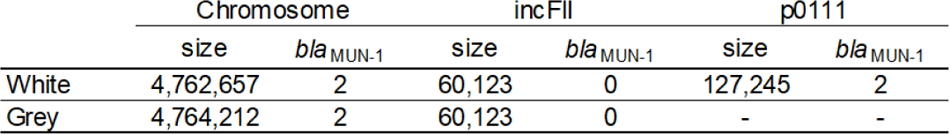
Differences in terms of contigs obtained after assembly using Unicycler between white and grey colonies after molecular characterization.

|  | Chromosome |  | incFII |  | p0111 |  |
| --- | --- | --- | --- | --- | --- | --- |
|  | size | *bla*<sub>MUN-1</sub> | size | *bla*<sub>MUN-1</sub> | size | *bla*<sub>MUN-1</sub> |
| White | 4,762,657 | 2 | 60,123 | 0 | 127,245 | 2 |
| Grey | 4,764,212 | 2 | 60,123 | 0 | - | - |

**Supplementary Table 2:**
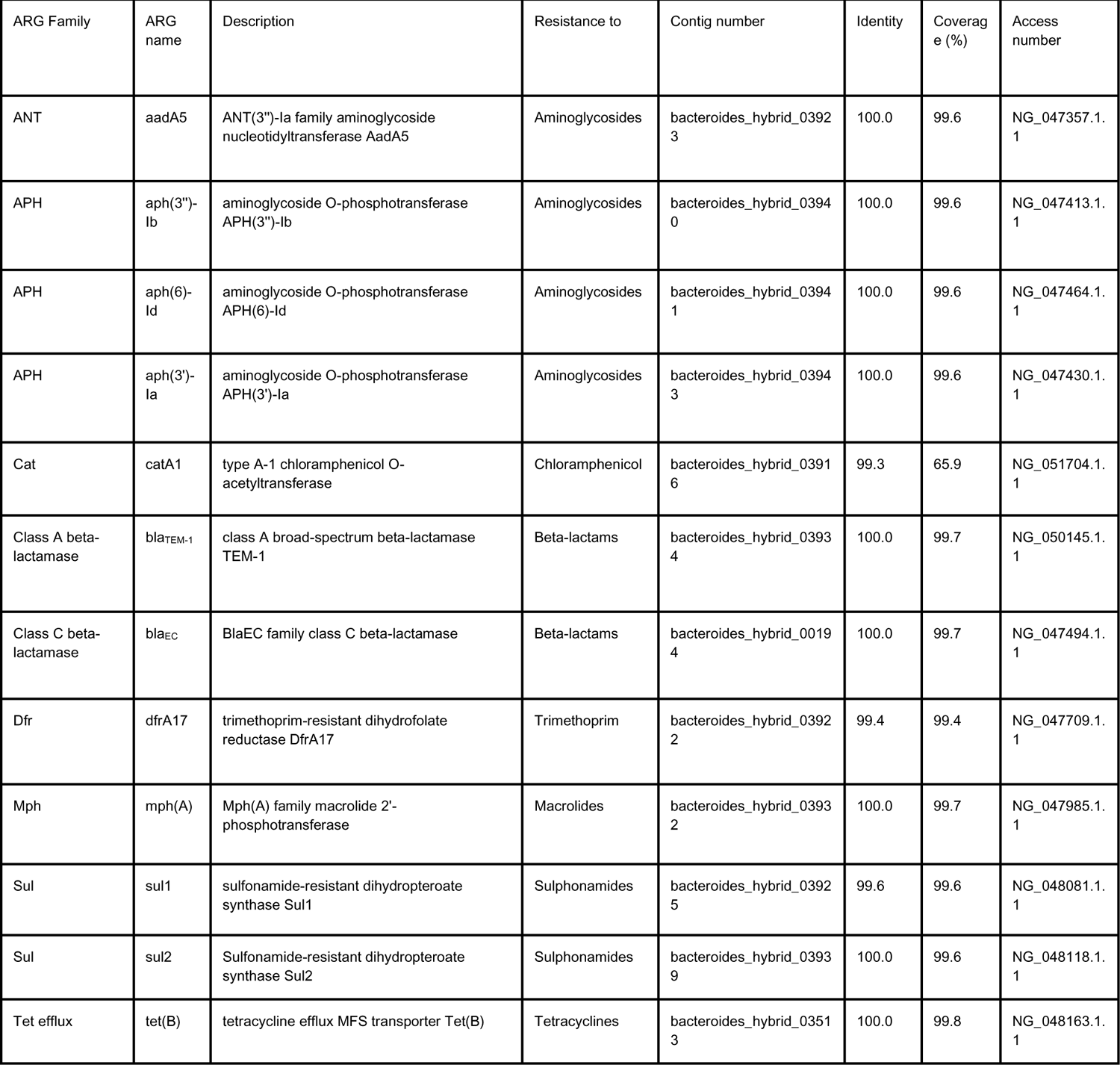
Antibiotic resistance genes (ARGs) found in the chromosome of the *E. coli* strain using ResFinder (v4.0).

**Supplementary Table 3:**
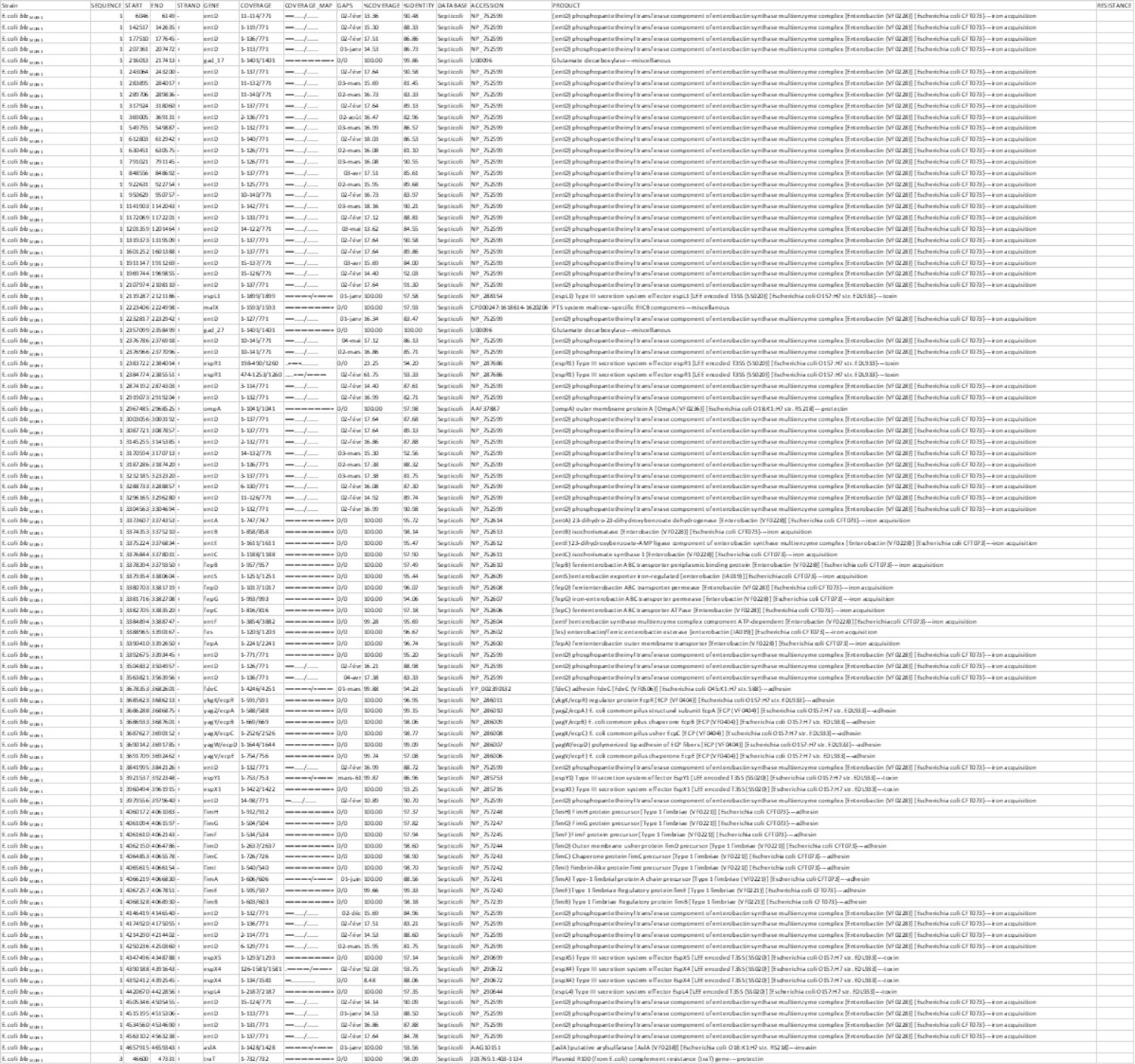
ABRicate analysis of the virulence gene found in the genome of the *E. coli* strain.

**Supplementary Table 4:**
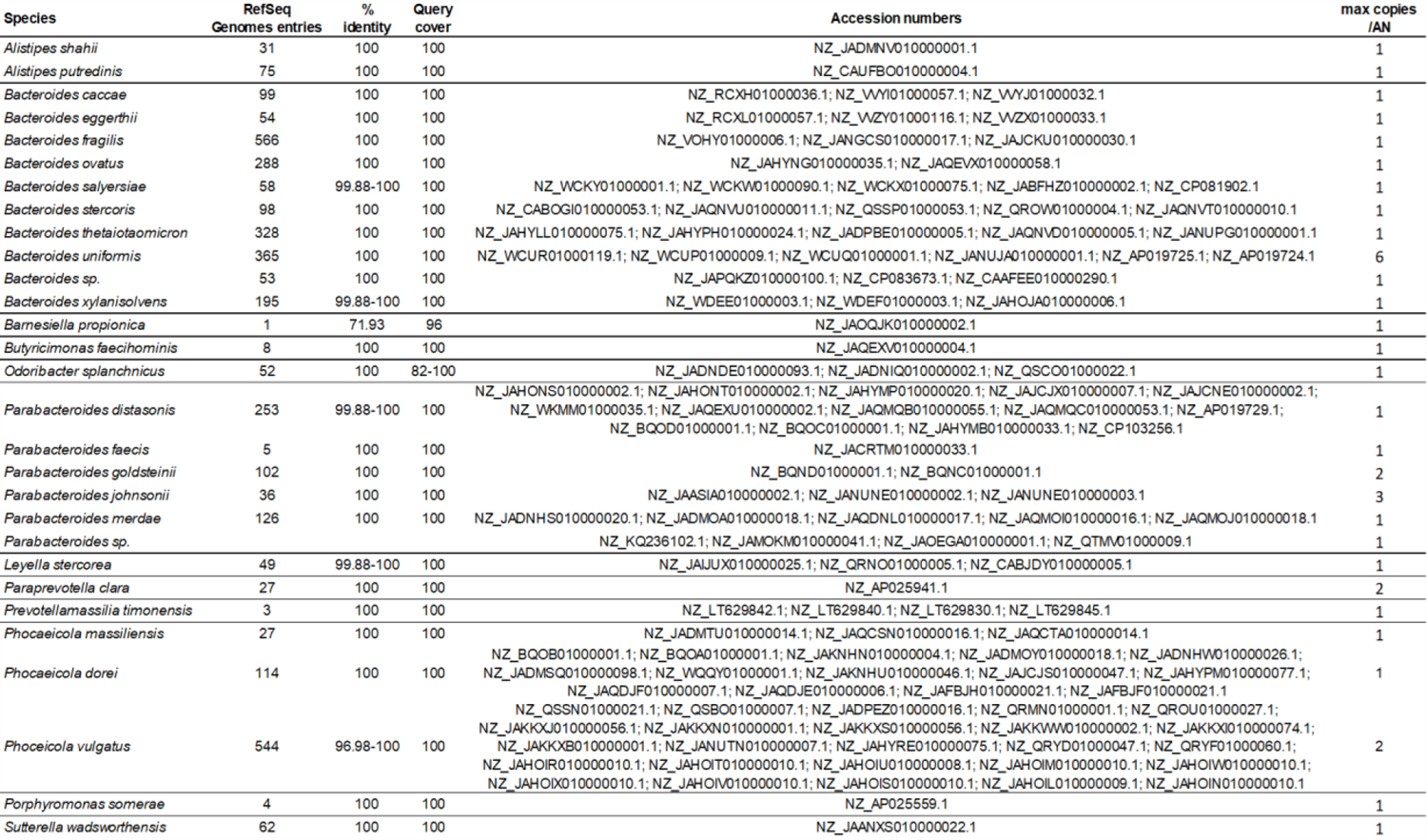
Hits obtained analyzing the *bla*_MUN-1_ gene distribution using BLASTN against RefSeq genomes database.

**Supplementary Table 5:**
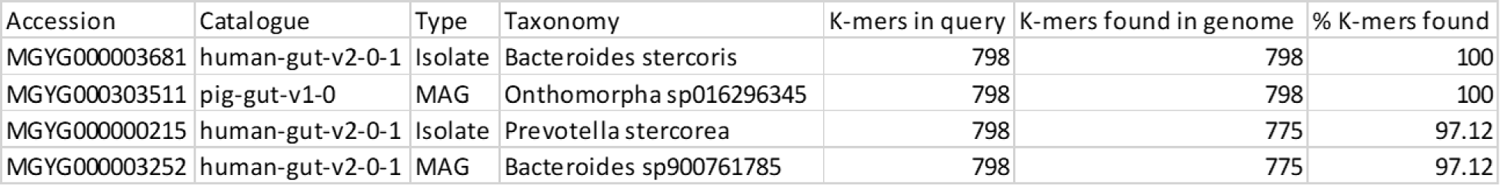
Hits obtained analyzing the *bla*_MUN-1_ gene distribution using MGnify.

**Supplementary Table 6:**
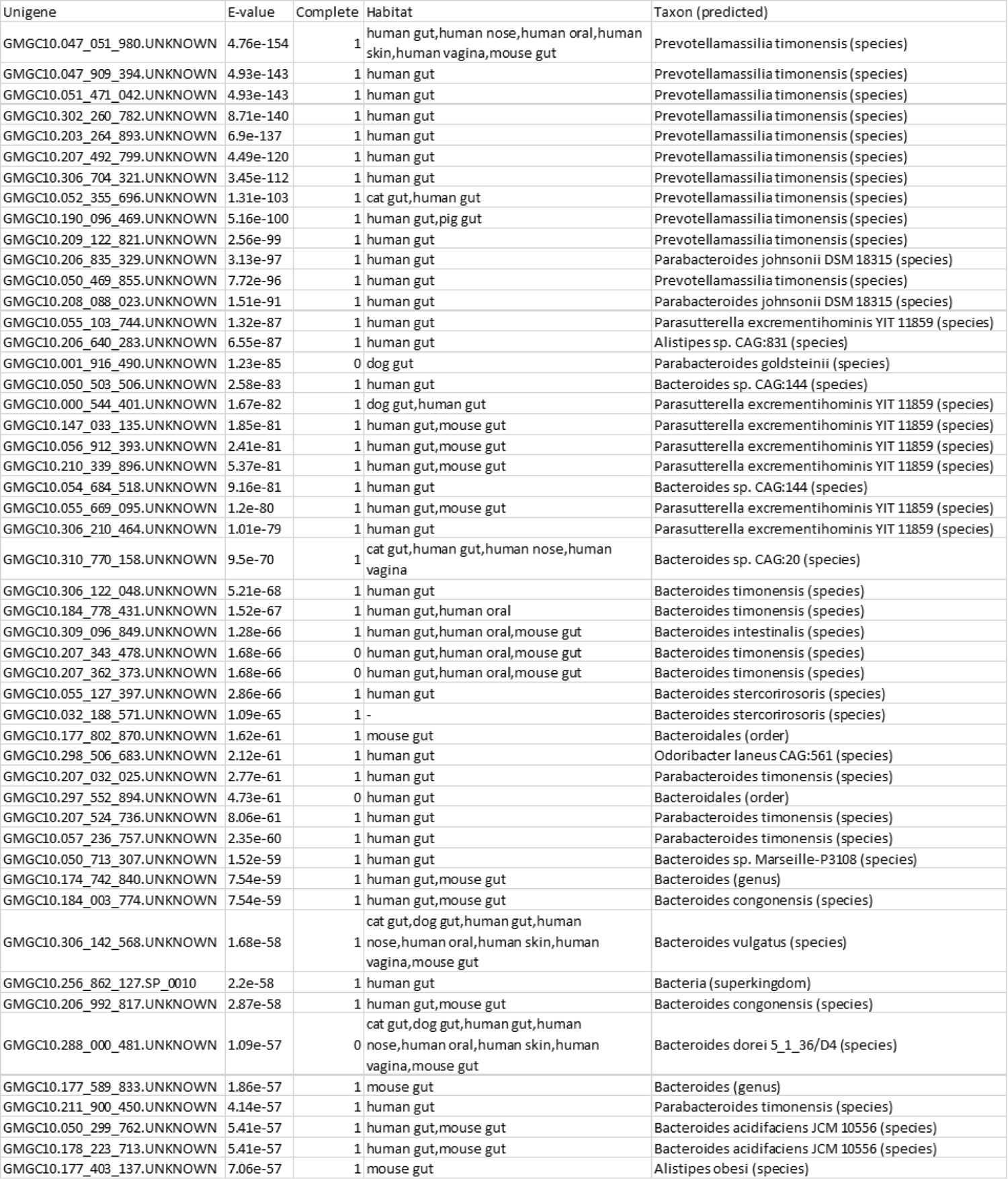
Hits obtained analyzing the MUN-1 protein distribution using the GMGC catalog.

**Supp Figure 1:**
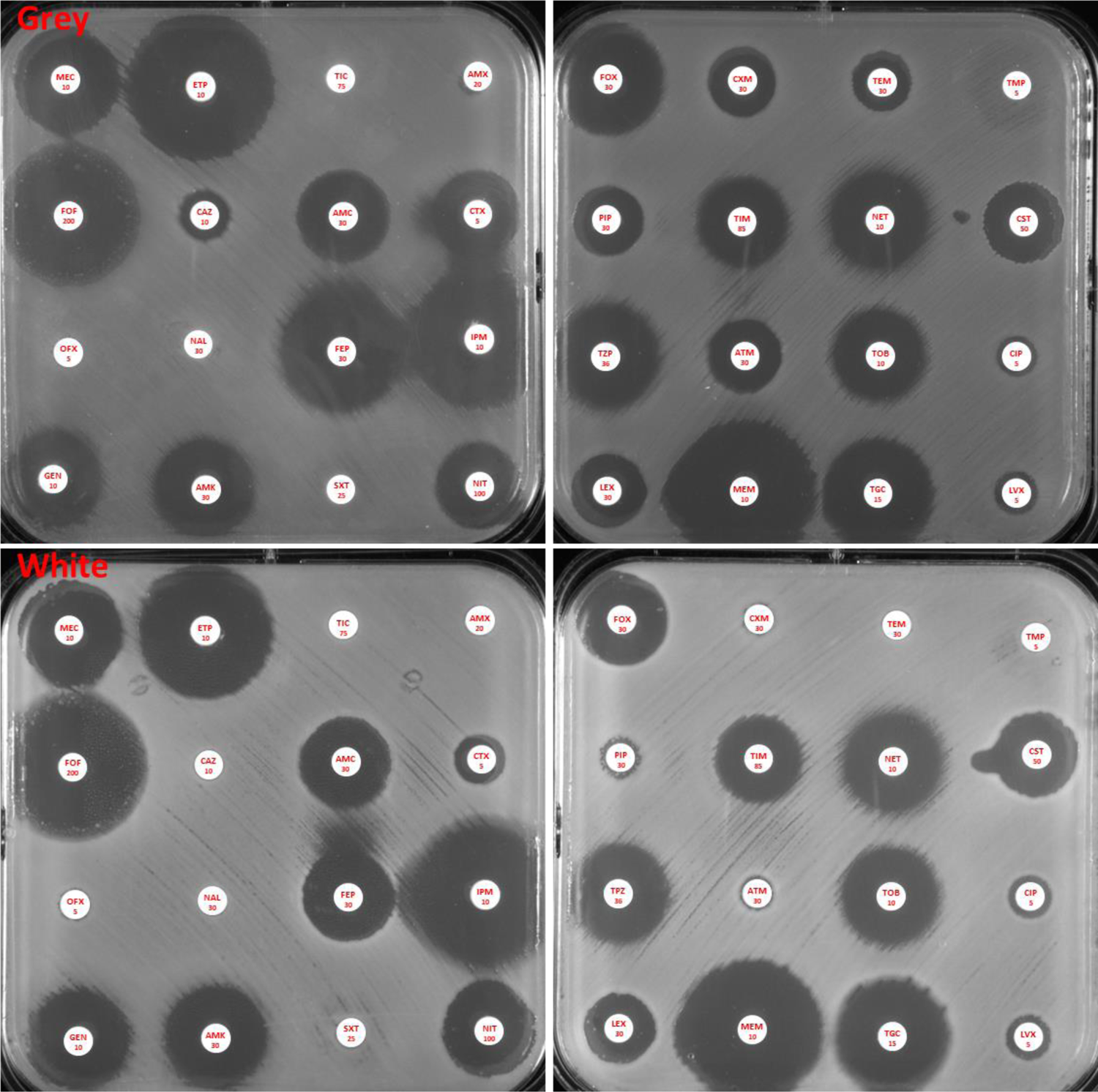
Antibiotic susceptibility testing. (MEC: mecillinam; ETP: ertapenem; TIC: ticarcillin; AMX: amoxicillin; FOF: fosfomycin; CAZ: ceftazidime; AMC: amoxicillin + clavulanic acid; CTX: cefotaxime; OFX: ofloxacin; NAL: nalidixic acid; FEP: cefepime; IPM: imipenem; GEN: gentamicin; AMK: amikacin; SXT: trimethoprim + sulfamethoxazole; NIT: nitrofurantoin; FOX: cefoxitin; CXM: cefuroxime; TEM: temocillin; TMP: trimethoprim; PIP: piperacillin; TIM: ticarcillin + clavulanic acid; NET: netilmicin; CST: colistin; TPZ: piperacillin + tazobactam; ATM: aztreonam; TOB: tobramycin; CIP: ciprofloxacin; LEX: cefalexin; MEM: meropenem; TGC: tigecycline; LVX: levofloxacin.

**Supplementary Figure 2:**
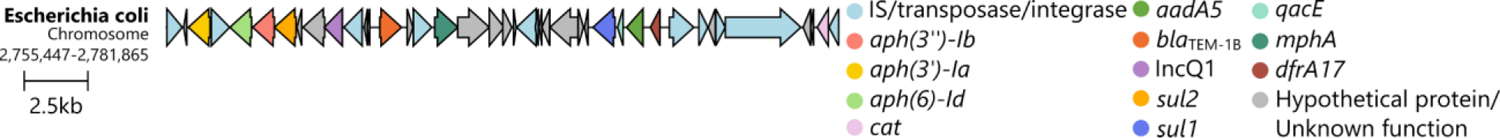
Antibiotic resistance genomic island found on the chromosome of the *E. coli* strain.

### Purification of the protein

Purification of the MUN-1 beta-lactamase was carried out by ion-exchange chromatography. *E. coli* TOP10 (pTOPO/*bla*_MUN-1_) was grown in 2 L of lysogeny broth (LB) containing 100 µg/mL of ampicillin, 50 µg/mL of kanamycin and 1 mM of IPTG. The overnight culture was centrifuged, and the obtained pellet was re-suspended in phosphate buffer pH 7 (0.1 M) and sonicated using a Vibra cellTM 75,186 sonicator (Thermo Fisher Scientific). After filtration using a 0.22 µm nitrocellulose filter, the crude extract was loaded in a prequilibrated SP-Sepharose column connected to an ÄktaPrime chromatography system (GE Healthcare) and eluted with a linear pH gradient using bis-Tris buffer at pH 6.4 (20 mM) and eluted with a linear NaCl gradient. The beta-lactamase recovered in the flow through was subsequently dialyzed against MES buffer at pH 6.4 (50 mM), loaded onto a Q-Sepharose column pre-equilibrated with the same buffer, and eluted with a linear NaCl gradient. The fractions containing the highest beta-lactamase activity, as determined by nitrocefin test (TRC Toronto, Canada), were pooled and dialyzed overnight against sodium phosphate buffer (pH 7, 0.1 M). The protein concentrations were measured using Bradford reagent (Sigma-Aldrich). The total protein content was measured by using a Bradford assay.

### Determination of beta-lactamase relative molecular mass

The relative purity of beta-lactamase was estimated by SDS-12% PAGE (GeneScript) analysis. Enzyme extracts were boiled for 10 min in a 1%SDS-3% beta-mercaptoethanol solution (GenScript) and then were subjected to electrophoresis with marker (GenScript) at room temperature.

### Enzymatic characterization

Purified beta-lactamase was used for kinetic measurements performed at room temperature in 100 mM sodium phosphate (pH 7.0). The initial rates of hydrolysis were determined with a Genesys 10S UV-visible spectrophotometer (Thermo Scientific). The following wavelengths and absorption coefficients were used: benzylpenicillin, 232 nm and ⋀ℇ of −1,100 M-1 cm-1; ampicillin, 240 nm and ⋀ℇ of −999 M-1 cm-1; ticarcillin, 235 nm and ⋀ℇ of −1,050 M-1 cm-1; piperacillin, 235 nm and ⋀ℇ of −1,070 M-1 cm-1; cephalothin, 262 nm and ⋀ℇ of −7,960 M-1 cm-1, cefoxitin, 265 nm and ⋀ℇ of −7,380 M-1 cm-1; ceftazidime, 260 nm and ⋀ℇ of −8,660 M-1 cm-1; cefepime, 264 nm and ⋀ℇ of −8240 M-1 cm-1, cefotaxime, 265 nm and ⋀ℇ of −6260 M-1 cm-1; imipenem, 297 nm and ⋀ℇ of −9210 M-1 cm-1; meropenem, 297 nm and ⋀ℇ of −9210 M-1 cm-1; aztreonam 318 nm and ⋀ℇ of −640 M-1 cm-1. The *Ki* values were determined by direct competition assays using 100 M nitrocefin. Inverse initial steady-state velocities (1/*V*0) were plotted against the inhibitor concentration ([I]) to obtain a straight line. The plots were linear and provided *y* intercept and slope values used for *Ki* determinations. *Ki* was determined by dividing the value for the *y* intercept by the slope of the line and then was corrected by taking into account the cephalothin affinity, using the following equation: *Ki* (corrected) = *Ki* (observed)/(1 [S]/*Km*), where [S] is the concentration of nitrocefin (100 µM) used in the assay and *Km* is the Michaelis constant determined for nitrocefin (45.5 µM).

### Susceptibility to beta-lactamase inhibitors

IC50 values were determined for clavulanic acid, tazobactam, and avibactam. Various concentrations of these inhibitors were pre-incubated with the purified enzyme MUN-1 3 min at room temperature, to determine the concentrations that reduced the hydrolysis rate with 100 M cephalothin by 50%. The results are expressed in nanomolar units.

## References

1. Ambler RP, Coulson AF, Frère JM, Ghuysen JM, Joris B, Forsman M, Levesque RC, Tiraby G, Waley SG. 1991. A standard numbering scheme for the class A beta-lactamases. Biochem J 276:269–270.

2. Ambler RP. 1980. The structure of beta-lactamases. Philos Trans R Soc Lond B Biol Sci 289:321– 331.

3. Ebmeyer S, Kristiansson E, Larsson DGJ. 2021. A framework for identifying the recent origins of mobile antibiotic resistance genes. Commun Biol 4:8.

4. Ghosh S, Sadowsky MJ, Roberts MC, Gralnick JA, LaPara TM. 2009. Sphingobacterium sp. strain PM2-P1-29 harbours a functional tet(X) gene encoding for the degradation of tetracycline. J Appl Microbiol 106:1336–1342.

5. Penders J, Stobberingh EE, Savelkoul PHM, Wolffs PFG. 2013. The human microbiome as a reservoir of antimicrobial resistance. Front Microbiol 4:87.

6. Ruppé E, Ghozlane A, Tap J, Pons N, Alvarez A-S, Maziers N, Cuesta T, Hernando-Amado S, Clares I, Martínez JL, Coque TM, Baquero F, Lanza VF, Máiz L, Goulenok T, de Lastours V, Amor N, Fantin B, Wieder I, Andremont A, van Schaik W, Rogers M, Zhang X, Willems RJL, de Brevern AG, Batto J-M, Blottière HM, Léonard P, Léjard V, Letur A, Levenez F, Weiszer K, Haimet F, Doré J, Kennedy SP, Ehrlich SD. 2019. Prediction of the intestinal resistome by a three-dimensional structure-based method. Nat Microbiol 4:112–123.

7. Sommer MOA, Dantas G, Church GM. 2009. Functional Characterization of the Antibiotic Resistance Reservoir in the Human Microflora. Science 325:1128–1131.

8. Kent AG, Vill AC, Shi Q, Satlin MJ, Brito IL. 2020. Widespread transfer of mobile antibiotic resistance genes within individual gut microbiomes revealed through bacterial Hi-C. Nat Commun 11:4379.

9. Petitjean M, Condamine B, Burdet C, Denamur E, Ruppé E. 2021. Phylum barrier and Escherichia coli intra-species phylogeny drive the acquisition of antibiotic-resistance genes. Microb Genom 7.

10. Zhou Z, Alikhan N-F, Mohamed K, Fan Y, Achtman M. 2020. The EnteroBase user’s guide, with case studies on Salmonella transmissions, Yersinia pestis phylogeny, and Escherichia core genomic diversity. Genome Res 30:138–152.

11. Gschwind R, Ugarcina Perovic S, Weiss M, Petitjean M, Lao J, Coelho LP, Ruppé E. 2023. ResFinderFG v2.0: a database of antibiotic resistance genes obtained by functional metagenomics. Nucleic Acids Res 51:W493–W500.

12. Mellmann A, Bletz S, Böking T, Kipp F, Becker K, Schultes A, Prior K, Harmsen D. 2016. Real-Time Genome Sequencing of Resistant Bacteria Provides Precision Infection Control in an Institutional Setting. J Clin Microbiol 54:2874–2881.

13. Philippon A, Jacquier H, Ruppé E, Labia R. 2019. Structure-based classification of class A beta-lactamases, an update. Curr Res Transl Med 67:115–122.

14. Altschul SF, Gish W, Miller W, Myers EW, Lipman DJ. 1990. Basic local alignment search tool. J Mol Biol 215:403–410.

15. Richardson L, Allen B, Baldi G, Beracochea M, Bileschi ML, Burdett T, Burgin J, Caballero-Pérez J, Cochrane G, Colwell LJ, Curtis T, Escobar-Zepeda A, Gurbich TA, Kale V, Korobeynikov A, Raj S, Rogers AB, Sakharova E, Sanchez S, Wilkinson DJ, Finn RD. 2023. MGnify: the microbiome sequence data analysis resource in 2023. Nucleic Acids Research 51:D753–D759.

16. Coelho LP, Alves R, Del Río ÁR, Myers PN, Cantalapiedra CP, Giner-Lamia J, Schmidt TS, Mende DR, Orakov A, Letunic I, Hildebrand F, Van Rossum T, Forslund SK, Khedkar S, Maistrenko OM, Pan S, Jia L, Ferretti P, Sunagawa S, Zhao X-M, Nielsen HB, Huerta-Cepas J, Bork P. 2022. Towards the biogeography of prokaryotic genes. Nature 601:252–256.

17. Kuch A, Zieniuk B, Żabicka D, Van de Velde S, Literacka E, Skoczyńska A, Hryniewicz W. 2020. Activity of temocillin against ESBL-, AmpC-, and/or KPC-producing Enterobacterales isolated in Poland. Eur J Clin Microbiol Infect Dis 39:1185–1191.

18. Rogers MB, Bennett TK, Payne CM, Smith CJ. 1994. Insertional activation of cepA leads to high-level beta-lactamase expression in Bacteroides fragilis clinical isolates. J Bacteriol 176:4376–4384.

19. Dupin C, Tamanai-Shacoori Z, Ehrmann E, Dupont A, Barloy-Hubler F, Bousarghin L, Bonnaure-Mallet M, Jolivet-Gougeon A. 2015. Oral Gram-negative anaerobic bacilli as a reservoir of β-lactam resistance genes facilitating infections with multiresistant bacteria. Int J Antimicrob Agents 45:99– 105.

20. Silva J, Aguilar C, Ayala G, Estrada MA, Garza-Ramos U, Lara-Lemus R, Ledezma L. 2000. TLA-1: a New Plasmid-Mediated Extended-Spectrum β-Lactamase from Escherichia coli. Antimicrobial Agents and Chemotherapy 44:997–1003.

21. Wang M, Jiang L, Wei J, Zhu H, Zhang J, Liu Z, Zhang W, He X, Liu Y, Li R, Xiao X, Sun Y, Zeng Z, Wang Z. 2022. Similarities of P1-Like Phage Plasmids and Their Role in the Dissemination of blaCTX-M-55. Microbiol Spectr 10:e0141022.

22. Pfeifer E, Moura de Sousa JA, Touchon M, Rocha EPC. 2021. Bacteria have numerous distinctive groups of phage–plasmids with conserved phage and variable plasmid gene repertoires. Nucleic Acids Research 49:2655–2673.

23. Rashtchian A, Booth SJ. 1981. Stability in Escherichia coli of an antibiotic resistance plasmid from Bacteroides fragilis. J Bacteriol 146:121–127.

24. Klümper U, Riber L, Dechesne A, Sannazzarro A, Hansen LH, Sørensen SJ, Smets BF. 2015. Broad host range plasmids can invade an unexpectedly diverse fraction of a soil bacterial community. ISME J 9:934–945.

25. Buchfink B, Xie C, Huson DH. 2015. Fast and sensitive protein alignment using DIAMOND. 1. Nat Methods 12:59–60.

26. Feldgarden M, Brover V, Haft DH, Prasad AB, Slotta DJ, Tolstoy I, Tyson GH, Zhao S, Hsu C-H, McDermott PF, Tadesse DA, Morales C, Simmons M, Tillman G, Wasilenko J, Folster JP, Klimke W. 2019. Validating the AMRFinder Tool and Resistance Gene Database by Using Antimicrobial Resistance Genotype-Phenotype Correlations in a Collection of Isolates. Antimicrob Agents Chemother 63:e00483–19.

27. Wick RR, Judd LM, Gorrie CL, Holt KE. 2017. Unicycler: Resolving bacterial genome assemblies from short and long sequencing reads. PLOS Computational Biology 13:e1005595.

28. Bortolaia V, Kaas RS, Ruppe E, Roberts MC, Schwarz S, Cattoir V, Philippon A, Allesoe RL, Rebelo AR, Florensa AF, Fagelhauer L, Chakraborty T, Neumann B, Werner G, Bender JK, Stingl K, Nguyen M, Coppens J, Xavier BB, Malhotra-Kumar S, Westh H, Pinholt M, Anjum MF, Duggett NA, Kempf I, Nykäsenoja S, Olkkola S, Wieczorek K, Amaro A, Clemente L, Mossong J, Losch S, Ragimbeau C, Lund O, Aarestrup FM. 2020. ResFinder 4.0 for predictions of phenotypes from genotypes. Journal of Antimicrobial Chemotherapy 75:3491–3500.

29. Carattoli A, Zankari E, García-Fernández A, Voldby Larsen M, Lund O, Villa L, Møller Aarestrup F, Hasman H. 2014. In silico detection and typing of plasmids using PlasmidFinder and plasmid multilocus sequence typing. Antimicrob Agents Chemother 58:3895–3903.

30. Schwengers O, Jelonek L, Dieckmann MA, Beyvers S, Blom J, Goesmann A. 2021. Bakta: rapid and standardized annotation of bacterial genomes via alignment-free sequence identification. Microb Genom 7:000685.

31. Gilchrist CLM, Chooi Y-H. 2021. clinker & clustermap.js: automatic generation of gene cluster comparison figures. Bioinformatics 37:2473–2475.

32. Kuraku S, Zmasek CM, Nishimura O, Katoh K. 2013. aLeaves facilitates on-demand exploration of metazoan gene family trees on MAFFT sequence alignment server with enhanced interactivity. Nucleic Acids Research 41:W22–W28.

33. Nguyen L-T, Schmidt HA, von Haeseler A, Minh BQ. 2015. IQ-TREE: A Fast and Effective Stochastic Algorithm for Estimating Maximum-Likelihood Phylogenies. Molecular Biology and Evolution 32:268–274.

34. Hoang DT, Chernomor O, von Haeseler A, Minh BQ, Vinh LS. 2018. UFBoot2: Improving the Ultrafast Bootstrap Approximation. Molecular Biology and Evolution 35:518–522.

