## Supplementary Table 3 for "Inter-phylum circulation of a beta-lactamase - encoding gene: a rare but observable event"

Supplementary Table 3: ABRicate analysis of the virulence gene found in the genome of the E. coli strain.

| Strain | SEQUENCE | START | END | STRAND | GENE | COVERAGE | COVERAGE MAP | GAPS | %COVERAGE | %IDENTITY | DATABASE | ACCESSION | PRODUCT | RESISTANCE |
|---|---|---|---|---|---|---|---|---|---|---|---|---|---|---|
| E. coli *bla* $_{MUN-1}$ | 1 | 6046 | 6149 | - | entD | 11-114/771 | ===...../...... | 02-févr | 13.36 | 90.48 | Septicoli | NP_752599 | (entD) phosphopantetheinyl transferase component of enterobactin synthase multienzyme complex [Enterobactin (VF0228)] [Escherichia coli CFT073]---iron acquisition | |
| E. coli *bla* $_{MUN-1}$ | 1 | 142517 | 142635 | + | entD | 1-119/771 | ===...../...... | 02-févr | 15.30 | 88.33 | Septicoli | NP_752599 | (entD) phosphopantetheinyl transferase component of enterobactin synthase multienzyme complex [Enterobactin (VF0228)] [Escherichia coli CFT073]---iron acquisition | |
| E. coli *bla* $_{MUN-1}$ | 1 | 177510 | 177645 | - | entD | 1-136/771 | ===...../...... | 02-févr | 17.51 | 86.86 | Septicoli | NP_752599 | (entD) phosphopantetheinyl transferase component of enterobactin synthase multienzyme complex [Enterobactin (VF0228)] [Escherichia coli CFT073]---iron acquisition | |
| E. coli *bla* $_{MUN-1}$ | 1 | 207361 | 207472 | + | entD | 1-113/771 | ===...../...... | 01-janv | 14.53 | 86.73 | Septicoli | NP_752599 | (entD) phosphopantetheinyl transferase component of enterobactin synthase multienzyme complex [Enterobactin (VF0228)] [Escherichia coli CFT073]---iron acquisition | |
| E. coli *bla* $_{MUN-1}$ | 1 | 216013 | 217413 | + | gad_17 | 1-1401/1401 | =============== | 0/0 | 100.00 | 99.86 | Septicoli | U00096 | Glutamate decarboxylase---miscellanous | |
| E. coli *bla* $_{MUN-1}$ | 1 | 243064 | 243200 | - | entD | 1-137/771 | ===...../...... | 02-févr | 17.64 | 90.58 | Septicoli | NP_752599 | (entD) phosphopantetheinyl transferase component of enterobactin synthase multienzyme complex [Enterobactin (VF0228)] [Escherichia coli CFT073]---iron acquisition | |
| E. coli *bla* $_{MUN-1}$ | 1 | 283895 | 284017 | + | entD | 11-132/771 | ===...../...... | 03-mars | 15.69 | 81.45 | Septicoli | NP_752599 | (entD) phosphopantetheinyl transferase component of enterobactin synthase multienzyme complex [Enterobactin (VF0228)] [Escherichia coli CFT073]---iron acquisition | |
| E. coli *bla* $_{MUN-1}$ | 1 | 289706 | 289836 | - | entD | 11-140/771 | ===...../...... | 02-mars | 16.73 | 83.33 | Septicoli | NP_752599 | (entD) phosphopantetheinyl transferase component of enterobactin synthase multienzyme complex [Enterobactin (VF0228)] [Escherichia coli CFT073]---iron acquisition | |
| E. coli *bla* $_{MUN-1}$ | 1 | 317924 | 318060 | + | entD | 1-137/771 | ===...../...... | 02-févr | 17.64 | 89.13 | Septicoli | NP_752599 | (entD) phosphopantetheinyl transferase component of enterobactin synthase multienzyme complex [Enterobactin (VF0228)] [Escherichia coli CFT073]---iron acquisition | |
| E. coli *bla* $_{MUN-1}$ | 1 | 369005 | 369131 | + | entD | 2-136/771 | ===...../...... | 02-août | 16.47 | 82.96 | Septicoli | NP_752599 | (entD) phosphopantetheinyl transferase component of enterobactin synthase multienzyme complex [Enterobactin (VF0228)] [Escherichia coli CFT073]---iron acquisition | |
| E. coli *bla* $_{MUN-1}$ | 1 | 549755 | 549887 | - | entD | 1-132/771 | ===...../...... | 03-mars | 16.99 | 86.57 | Septicoli | NP_752599 | (entD) phosphopantetheinyl transferase component of enterobactin synthase multienzyme complex [Enterobactin (VF0228)] [Escherichia coli CFT073]---iron acquisition | |
| E. coli *bla* $_{MUN-1}$ | 1 | 612803 | 612942 | + | entD | 1-140/771 | ===...../...... | 02-févr | 18.03 | 86.53 | Septicoli | NP_752599 | (entD) phosphopantetheinyl transferase component of enterobactin synthase multienzyme complex [Enterobactin (VF0228)] [Escherichia coli CFT073]---iron acquisition | |
| E. coli *bla* $_{MUN-1}$ | 1 | 630451 | 630575 | - | entD | 1-126/771 | ===...../...... | 02-mars | 16.08 | 81.10 | Septicoli | NP_752599 | (entD) phosphopantetheinyl transferase component of enterobactin synthase multienzyme complex [Enterobactin (VF0228)] [Escherichia coli CFT073]---iron acquisition | |
| E. coli *bla* $_{MUN-1}$ | 1 | 791021 | 791145 | - | entD | 1-126/771 | ===...../...... | 03-mars | 16.08 | 90.55 | Septicoli | NP_752599 | (entD) phosphopantetheinyl transferase component of enterobactin synthase multienzyme complex [Enterobactin (VF0228)] [Escherichia coli CFT073]---iron acquisition | |
| E. coli *bla* $_{MUN-1}$ | 1 | 848556 | 848692 | - | entD | 1-137/771 | ===...../...... | 03-avr | 17.51 | 85.61 | Septicoli | NP_752599 | (entD) phosphopantetheinyl transferase component of enterobactin synthase multienzyme complex [Enterobactin (VF0228)] [Escherichia coli CFT073]---iron acquisition | |
| E. coli *bla* $_{MUN-1}$ | 1 | 922631 | 922754 | + | entD | 1-125/771 | ===...../...... | 02-mars | 15.95 | 89.68 | Septicoli | NP_752599 | (entD) phosphopantetheinyl transferase component of enterobactin synthase multienzyme complex [Enterobactin (VF0228)] [Escherichia coli CFT073]---iron acquisition | |
| E. coli *bla* $_{MUN-1}$ | 1 | 950629 | 950757 | - | entD | 10-140/771 | ===...../...... | 02-févr | 16.73 | 83.97 | Septicoli | NP_752599 | (entD) phosphopantetheinyl transferase component of enterobactin synthase multienzyme complex [Enterobactin (VF0228)] [Escherichia coli CFT073]---iron acquisition | |
| E. coli *bla* $_{MUN-1}$ | 1 | 1E+06 | 1E+06 | + | entD | 1-142/771 | ===...../...... | 03-mars | 18.16 | 90.21 | Septicoli | NP_752599 | (entD) phosphopantetheinyl transferase component of enterobactin synthase multienzyme complex [Enterobactin (VF0228)] [Escherichia coli CFT073]---iron acquisition | |
| E. coli *bla* $_{MUN-1}$ | 1 | 1E+06 | 1E+06 | + | entD | 1-133/771 | ===...../...... | 02-févr | 17.12 | 88.81 | Septicoli | NP_752599 | (entD) phosphopantetheinyl transferase component of enterobactin synthase multienzyme complex [Enterobactin (VF0228)] [Escherichia coli CFT073]---iron acquisition | |
| E. coli *bla* $_{MUN-1}$ | 1 | 1E+06 | 1E+06 | + | entD | 14-122/771 | ===...../...... | 03-mai | 13.62 | 84.55 | Septicoli | NP_752599 | (entD) phosphopantetheinyl transferase component of enterobactin synthase multienzyme complex [Enterobactin (VF0228)] [Escherichia coli CFT073]---iron acquisition | |
| E. coli *bla* $_{MUN-1}$ | 1 | 1E+06 | 1E+06 | + | entD | 1-137/771 | ===...../...... | 02-févr | 17.64 | 90.58 | Septicoli | NP_752599 | (entD) phosphopantetheinyl transferase component of enterobactin synthase multienzyme complex [Enterobactin (VF0228)] [Escherichia coli CFT073]---iron acquisition | |
| E. coli *bla* $_{MUN-1}$ | 1 | 2E+06 | 2E+06 | + | entD | 1-137/771 | ===...../...... | 02-févr | 17.64 | 89.86 | Septicoli | NP_752599 | (entD) phosphopantetheinyl transferase component of enterobactin synthase multienzyme complex [Enterobactin (VF0228)] [Escherichia coli CFT073]---iron acquisition | |
| E. coli *bla* $_{MUN-1}$ | 1 | 2E+06 | 2E+06 | - | entD | 15-137/771 | ===...../...... | 03-avr | 15.69 | 84.00 | Septicoli | NP_752599 | (entD) phosphopantetheinyl transferase component of enterobactin synthase multienzyme complex [Enterobactin (VF0228)] [Escherichia coli CFT073]---iron acquisition | |
| E. coli *bla* $_{MUN-1}$ | 1 | 2E+06 | 2E+06 | - | entD | 15-126/771 | ===...../...... | 02-févr | 14.40 | 92.03 | Septicoli | NP_752599 | (entD) phosphopantetheinyl transferase component of enterobactin synthase multienzyme complex [Enterobactin (VF0228)] [Escherichia coli CFT073]---iron acquisition | |
| E. coli *bla* $_{MUN-1}$ | 1 | 2E+06 | 2E+06 | - | entD | 1-137/771 | ===...../...... | 02-févr | 17.64 | 91.30 | Septicoli | NP_752599 | (entD) phosphopantetheinyl transferase component of enterobactin synthase multienzyme complex [Enterobactin (VF0228)] [Escherichia coli CFT073]---iron acquisition | |
| E. coli *bla* $_{MUN-1}$ | 1 | 2E+06 | 2E+06 | - | espL1 | 1-1899/1899 | ========/====== | 01-janv | 100.00 | 97.58 | Septicoli | NP_288154 | (espL1) Type III secretion system effector espL1 [LEE encoded T3SS (SS020)] [Escherichia coli O157:H7 str. EDL933]---toxin | |
| E. coli *bla* $_{MUN-1}$ | 1 | 2E+06 | 2E+06 | - | malX | 1-1593/1593 | =============== | 0/0 | 100.00 | 97.93 | Septicoli | CP000247:1618614-1620206 | PTS system maltose-specific EIICB component---miscellanous | |
| E. coli *bla* $_{MUN-1}$ | 1 | 2E+06 | 2E+06 | + | entD | 1-127/771 | ===...../...... | 01-janv | 16.34 | 83.47 | Septicoli | NP_752599 | (entD) phosphopantetheinyl transferase component of enterobactin synthase multienzyme complex [Enterobactin (VF0228)] [Escherichia coli CFT073]---iron acquisition | |
| E. coli *bla* $_{MUN-1}$ | 1 | 2E+06 | 2E+06 | + | gad_27 | 1-1401/1401 | =============== | 0/0 | 100.00 | 100.00 | Septicoli | U00096 | Glutamate decarboxylase---miscellanous | |
| E. coli *bla* $_{MUN-1}$ | 1 | 2E+06 | 2E+06 | - | entD | 10-145/771 | ===...../...... | 04-mai | 17.12 | 86.13 | Septicoli | NP_752599 | (entD) phosphopantetheinyl transferase component of enterobactin synthase multienzyme complex [Enterobactin (VF0228)] [Escherichia coli CFT073]---iron acquisition | |
| E. coli *bla* $_{MUN-1}$ | 1 | 2E+06 | 2E+06 | - | entD | 10-141/771 | ===...../...... | 02-mars | 16.86 | 85.71 | Septicoli | NP_752599 | (entD) phosphopantetheinyl transferase component of enterobactin synthase multienzyme complex [Enterobactin (VF0228)] [Escherichia coli CFT073]---iron acquisition | |
| E. coli *bla* $_{MUN-1}$ | 1 | 2E+06 | 2E+06 | + | espR1 | 198-490/1260 | ..====......... | 0/0 | 23.25 | 94.20 | Septicoli | NP_287686 | (espR1) Type III secretion system effector espR1 [LEE encoded T3SS (SS020)] [Escherichia coli O157:H7 str. EDL933]---toxin | |
| E. coli *bla* $_{MUN-1}$ | 1 | 2E+06 | 2E+06 | + | espR1 | 474-1253/1260 | .....===/====== | 02-févr | 61.75 | 93.33 | Septicoli | NP_287686 | (espR1) Type III secretion system effector espR1 [LEE encoded T3SS (SS020)] [Escherichia coli O157:H7 str. EDL933]---toxin | |
| E. coli *bla* $_{MUN-1}$ | 1 | 3E+06 | 3E+06 | + | entD | 3-114/771 | ===...../...... | 02-févr | 14.40 | 87.61 | Septicoli | NP_752599 | (entD) phosphopantetheinyl transferase component of enterobactin synthase multienzyme complex [Enterobactin (VF0228)] [Escherichia coli CFT073]---iron acquisition | |
| E. coli *bla* $_{MUN-1}$ | 1 | 3E+06 | 3E+06 | + | entD | 1-132/771 | ===...../...... | 02-févr | 16.99 | 82.71 | Septicoli | NP_752599 | (entD) phosphopantetheinyl transferase component of enterobactin synthase multienzyme complex [Enterobactin (VF0228)] [Escherichia coli CFT073]---iron acquisition | |
| E. coli *bla* $_{MUN-1}$ | 1 | 3E+06 | 3E+06 | + | ompA | 1-1041/1041 | =============== | 0/0 | 100.00 | 97.98 | Septicoli | AAF37887 | (ompA) outer membrane protein A [OmpA (VF0236)] [Escherichia coli O18:K1:H7 str. RS218]---protectin | |
| E. coli *bla* $_{MUN-1}$ | 1 | 3E+06 | 3E+06 | - | entD | 1-137/771 | ===...../...... | 02-févr | 17.64 | 87.68 | Septicoli | NP_752599 | (entD) phosphopantetheinyl transferase component of enterobactin synthase multienzyme complex [Enterobactin (VF0228)] [Escherichia coli CFT073]---iron acquisition | |
| E. coli *bla* $_{MUN-1}$ | 1 | 3E+06 | 3E+06 | - | entD | 1-137/771 | ===...../...... | 02-févr | 17.64 | 89.13 | Septicoli | NP_752599 | (entD) phosphopantetheinyl transferase component of enterobactin synthase multienzyme complex [Enterobactin (VF0228)] [Escherichia coli CFT073]---iron acquisition | |
| E. coli *bla* $_{MUN-1}$ | 1 | 3E+06 | 3E+06 | + | entD | 2-132/771 | ===...../...... | 02-févr | 16.86 | 87.88 | Septicoli | NP_752599 | (entD) phosphopantetheinyl transferase component of enterobactin synthase multienzyme complex [Enterobactin (VF0228)] [Escherichia coli CFT073]---iron acquisition | |
| E. coli *bla* $_{MUN-1}$ | 1 | 3E+06 | 3E+06 | + | entD | 14-132/771 | ===...../...... | 03-mars | 15.30 | 92.56 | Septicoli | NP_752599 | (entD) phosphopantetheinyl transferase component of enterobactin synthase multienzyme complex [Enterobactin (VF0228)] [Escherichia coli CFT073]---iron acquisition | |
| E. coli *bla* $_{MUN-1}$ | 1 | 3E+06 | 3E+06 | + | entD | 1-136/771 | ===...../...... | 02-mars | 17.38 | 88.32 | Septicoli | NP_752599 | (entD) phosphopantetheinyl transferase component of enterobactin synthase multienzyme complex [Enterobactin (VF0228)] [Escherichia coli CFT073]---iron acquisition | |
| E. coli *bla* $_{MUN-1}$ | 1 | 3E+06 | 3E+06 | - | entD | 3-137/771 | ===...../...... | 03-mars | 17.38 | 81.75 | Septicoli | NP_752599 | (entD) phosphopantetheinyl transferase component of enterobactin synthase multienzyme complex [Enterobactin (VF0228)] [Escherichia coli CFT073]---iron acquisition | |
| E. coli *bla* $_{MUN-1}$ | 1 | 3E+06 | 3E+06 | + | entD | 6-130/771 | ===...../...... | 02-févr | 16.08 | 87.30 | Septicoli | NP_752599 | (entD) phosphopantetheinyl transferase component of enterobactin synthase multienzyme complex [Enterobactin (VF0228)] [Escherichia coli CFT073]---iron acquisition | |
| E. coli *bla* $_{MUN-1}$ | 1 | 3E+06 | 3E+06 | + | entD | 11-126/771 | ===...../...... | 02-févr | 14.92 | 89.74 | Septicoli | NP_752599 | (entD) phosphopantetheinyl transferase component of enterobactin synthase multienzyme complex [Enterobactin (VF0228)] [Escherichia coli CFT073]---iron acquisition | |
| E. coli *bla* $_{MUN-1}$ | 1 | 3E+06 | 3E+06 | - | entD | 1-132/771 | ===...../...... | 02-févr | 16.99 | 90.98 | Septicoli | NP_752599 | (entD) phosphopantetheinyl transferase component of enterobactin synthase multienzyme complex [Enterobactin (VF0228)] [Escherichia coli CFT073]---iron acquisition | |
| E. coli *bla* $_{MUN-1}$ | 1 | 3E+06 | 3E+06 | - | entA | 1-747/747 | =============== | 0/0 | 100.00 | 95.72 | Septicoli | NP_752614 | (entA) 23-dihydro-23-dihydroxybenzoate dehydrogenase [Enterobactin (VF0228)] [Escherichia coli CFT073]---iron acquisition | |
| E. coli *bla* $_{MUN-1}$ | 1 | 3E+06 | 3E+06 | - | entB | 1-858/858 | =============== | 0/0 | 100.00 | 98.14 | Septicoli | NP_752613 | (entB) isochorismatase [Enterobactin (VF0228)] [Escherichia coli CFT073]---iron acquisition | |
| E. coli *bla* $_{MUN-1}$ | 1 | 3E+06 | 3E+06 | - | entE | 1-1611/1611 | =============== | 0/0 | 100.00 | 95.47 | Septicoli | NP_752612 | (entE) 23-dihydroxybenzoate-AMP ligase component of enterobactin synthase multienzyme complex [Enterobactin (VF0228)] [Escherichia coli CFT073]---iron acquisition | |
| E. coli *bla* $_{MUN-1}$ | 1 | 3E+06 | 3E+06 | - | entC | 1-1188/1188 | =============== | 0/0 | 100.00 | 97.90 | Septicoli | NP_752611 | (entC) isochorismate synthase 1 [Enterobactin (VF0228)] [Escherichia coli CFT073]---iron acquisition | |
| E. coli *bla* $_{MUN-1}$ | 1 | 3E+06 | 3E+06 | + | fepB | 1-957/957 | =============== | 0/0 | 100.00 | 97.49 | Septicoli | NP_752610 | (fepB) ferrienterobactin ABC transporter periplasmic binding protein [Enterobactin (VF0228)] [Escherichia coli CFT073]---iron acquisition | |
| E. coli *bla* $_{MUN-1}$ | 1 | 3E+06 | 3E+06 | - | entS | 1-1251/1251 | =============== | 0/0 | 100.00 | 95.44 | Septicoli | NP_752609 | (entS) enterobactin exporter iron-regulated [enterobactin (IA019)] [Escherichia coli CFT073]---iron acquisition | |
| E. coli *bla* $_{MUN-1}$ | 1 | 3E+06 | 3E+06 | + | fepD | 1-1017/1017 | =============== | 0/0 | 100.00 | 96.07 | Septicoli | NP_752608 | (fepD) ferrienterobactin ABC transporter permease [Enterobactin (VF0228)] [Escherichia coli CFT073]---iron acquisition | |
| E. coli *bla* $_{MUN-1}$ | 1 | 3E+06 | 3E+06 | + | fepG | 1-993/993 | =============== | 0/0 | 100.00 | 94.06 | Septicoli | NP_752607 | (fepG) iron-enterobactin ABC transporter permease [Enterobactin (VF0228)] [Escherichia coli CFT073]---iron acquisition | |
| E. coli *bla* $_{MUN-1}$ | 1 | 3E+06 | 3E+06 | + | fepC | 1-816/816 | =============== | 0/0 | 100.00 | 97.18 | Septicoli | NP_752606 | (fepC) ferrienterobactin ABC transporter ATPase [Enterobactin (VF0228)] [Escherichia coli CFT073]---iron acquisition | |
| E. coli *bla* $_{MUN-1}$ | 1 | 3E+06 | 3E+06 | - | entF | 1-3854/3882 | =============== | 0/0 | 99.28 | 95.69 | Septicoli | NP_752604 | (entF) enterobactin synthase multienzyme complex component ATP-dependent [Enterobactin (VF0228)] [Escherichia coli CFT073]---iron acquisition | |
| E. coli *bla* $_{MUN-1}$ | 1 | 3E+06 | 3E+06 | - | fes | 1-1203/1203 | =============== | 0/0 | 100.00 | 96.67 | Septicoli | NP_752602 | (fes) enterobactin/ferric enterobactin esterase [enterobactin (IA019)] [Escherichia coli CFT073]---iron acquisition | |
| E. coli *bla* $_{MUN-1}$ | 1 | 3E+06 | 3E+06 | + | fepA | 1-2241/2241 | =============== | 0/0 | 100.00 | 96.74 | Septicoli | NP_752600 | (fepA) ferrienterobactin outer membrane transporter [Enterobactin (VF0228)] [Escherichia coli CFT073]---iron acquisition | |
| E. coli *bla* $_{MUN-1}$ | 1 | 3E+06 | 3E+06 | + | entD | 1-771/771 | =============== | 0/0 | 100.00 | 95.20 | Septicoli | NP_752599 | (entD) phosphopantetheinyl transferase component of enterobactin synthase multienzyme complex [Enterobactin (VF0228)] [Escherichia coli CFT073]---iron acquisition | |
| E. coli *bla* $_{MUN-1}$ | 1 | 4E+06 | 4E+06 | - | entD | 1-126/771 | ===...../...... | 02-févr | 16.21 | 88.98 | Septicoli | NP_752599 | (entD) phosphopantetheinyl transferase component of enterobactin synthase multienzyme complex [Enterobactin (VF0228)] [Escherichia coli CFT073]---iron acquisition | |
| E. coli *bla* $_{MUN-1}$ | 1 | 4E+06 | 4E+06 | + | entD | 1-136/771 | ===...../...... | 04-avr | 17.38 | 83.33 | Septicoli | NP_752599 | (entD) phosphopantetheinyl transferase component of enterobactin synthase multienzyme complex [Enterobactin (VF0228)] [Escherichia coli CFT073]---iron acquisition | |
| E. coli *bla* $_{MUN-1}$ | 1 | 4E+06 | 4E+06 | - | fdeC | 1-4246/4251 | ========/====== | 01-mars | 99.88 | 94.23 | Septicoli | YP_002390132 | (fdeC) adhesin FdeC [FdeC (VF0506)] [Escherichia coli O45:K1:H7 str. S88]---adhesin | |
| E. coli *bla* $_{MUN-1}$ | 1 | 4E+06 | 4E+06 | + | ykgK/ecpR | 1-591/591 | =============== | 0/0 | 100.00 | 96.95 | Septicoli | NP_286011 | (ykgK/ecpR) regulator protein EcpR [ECP (VF0404)] [Escherichia coli O157:H7 str. EDL933]---adhesin | |
| E. coli *bla* $_{MUN-1}$ | 1 | 4E+06 | 4E+06 | + | yagZ/ecpA | 1-588/588 | =============== | 0/0 | 100.00 | 99.15 | Septicoli | NP_286010 | (yagZ/ecpA) E. coli common pilus structural subunit EcpA [ECP (VF0404)] [Escherichia coli O157:H7 str. EDL933]---adhesin | |
| E. coli *bla* $_{MUN-1}$ | 1 | 4E+06 | 4E+06 | + | yagY/ecpB | 1-669/669 | =============== | 0/0 | 100.00 | 98.06 | Septicoli | NP_286009 | (yagY/ecpB) E. coli common pilus chaperone EcpB [ECP (VF0404)] [Escherichia coli O157:H7 str. EDL933]---adhesin | |
| E. coli *bla* $_{MUN-1}$ | 1 | 4E+06 | 4E+06 | + | yagX/ecpC | 1-2526/2526 | =============== | 0/0 | 100.00 | 98.77 | Septicoli | NP_286008 | (yagX/ecpC) E. coli common pilus usher EcpC [ECP (VF0404)] [Escherichia coli O157:H7 str. EDL933]---adhesin | |
| E. coli *bla* $_{MUN-1}$ | 1 | 4E+06 | 4E+06 | + | yagW/ecpD | 1-1644/1644 | =============== | 0/0 | 100.00 | 99.09 | Septicoli | NP_286007 | (yagW/ecpD) polymerized tip adhesin of ECP fibers [ECP (VF0404)] [Escherichia coli O157:H7 str. EDL933]---adhesin | |
| E. coli *bla* $_{MUN-1}$ | 1 | 4E+06 | 4E+06 | + | yagV/ecpE | 1-754/756 | =============== | 0/0 | 99.74 | 97.08 | Septicoli | NP_286006 | (yagV/ecpE) E. coli common pilus chaperone EcpE [ECP (VF0404)] [Escherichia coli O157:H7 str. EDL933]---adhesin | |
| E. coli *bla* $_{MUN-1}$ | 1 | 4E+06 | 4E+06 | + | entD | 1-132/771 | ===...../...... | 02-févr | 16.99 | 88.72 | Septicoli | NP_752599 | (entD) phosphopantetheinyl transferase component of enterobactin synthase multienzyme complex [Enterobactin (VF0228)] [Escherichia coli CFT073]---iron acquisition | |
| E. coli *bla* $_{MUN-1}$ | 1 | 4E+06 | 4E+06 | - | espY1 | 1-753/753 | ========/====== | mars-61 | 99.87 | 86.96 | Septicoli | NP_285753 | (espY1) Type III secretion system effector EspY1 [LEE encoded T3SS (SS020)] [Escherichia coli O157:H7 str. EDL933]---toxin | |
| E. coli *bla* $_{MUN-1}$ | 1 | 4E+06 | 4E+06 | + | espX1 | 1-1422/1422 | =============== | 0/0 | 100.00 | 93.25 | Septicoli | NP_285716 | (espX1) Type III secretion system effector EspX1 [LEE encoded T3SS (SS020)] [Escherichia coli O157:H7 str. EDL933]---toxin | |
| E. coli *bla* $_{MUN-1}$ | 1 | 4E+06 | 4E+06 | + | entD | 14-98/771 | ==....../...... | 02-févr | 10.89 | 90.70 | Septicoli | NP_752599 | (entD) phosphopantetheinyl transferase component of enterobactin synthase multienzyme complex [Enterobactin (VF0228)] [Escherichia coli CFT073]---iron acquisition | |
| E. coli *bla* $_{MUN-1}$ | 1 | 4E+06 | 4E+06 | - | fimH | 1-912/912 | =============== | 0/0 | 100.00 | 97.37 | Septicoli | NP_757248 | (fimH) FimH protein precursor [Type 1 fimbriae (VF0221)] [Escherichia coli CFT073]---adhesin | |
| E. coli *bla* $_{MUN-1}$ | 1 | 4E+06 | 4E+06 | - | fimG | 1-504/504 | =============== | 0/0 | 100.00 | 97.82 | Septicoli | NP_757247 | (fimG) FimG protein precursor [Type 1 fimbriae (VF0221)] [Escherichia coli CFT073]---adhesin | |
| E. coli *bla* $_{MUN-1}$ | 1 | 4E+06 | 4E+06 | - | fimF | 1-534/534 | =============== | 0/0 | 100.00 | 97.94 | Septicoli | NP_757245 | (fimF) FimF protein precursor [Type 1 fimbriae (VF0221)] [Escherichia coli CFT073]---adhesin | |
| E. coli *bla* $_{MUN-1}$ | 1 | 4E+06 | 4E+06 | - | fimD | 1-2637/2637 | =============== | 0/0 | 100.00 | 98.60 | Septicoli | NP_757244 | (fimD) Outer membrane usher protein fimD precursor [Type 1 fimbriae (VF0221)] [Escherichia coli CFT073]---adhesin | |
| E. coli *bla* $_{MUN-1}$ | 1 | 4E+06 | 4E+06 | - | fimC | 1-726/726 | =============== | 0/0 | 100.00 | 98.90 | Septicoli | NP_757243 | (fimC) Chaperone protein fimC precursor [Type 1 fimbriae (VF0221)] [Escherichia coli CFT073]---adhesin | |
| E. coli *bla* $_{MUN-1}$ | 1 | 4E+06 | 4E+06 | - | fimI | 1-540/540 | =============== | 0/0 | 100.00 | 98.70 | Septicoli | NP_757242 | (fimI) Fimbrin-like protein fimI precursor [Type 1 fimbriae (VF0221)] [Escherichia coli CFT073]---adhesin | |
| E. coli *bla* $_{MUN-1}$ | 1 | 4E+06 | 4E+06 | - | fimA | 1-606/606 | ========/====== | 01-juin | 100.00 | 88.56 | Septicoli | NP_757241 | (fimA) Type-1 fimbrial protein A chain precursor [Type 1 fimbriae (VF0221)] [Escherichia coli CFT073]---adhesin | |
| E. coli *bla* $_{MUN-1}$ | 1 | 4E+06 | 4E+06 | - | fimE | 1-595/597 | =============== | 0/0 | 99.66 | 99.33 | Septicoli | NP_757240 | (fimE) Type 1 fimbriae Regulatory protein fimE [Type 1 fimbriae (VF0221)] [Escherichia coli CFT073]---adhesin | |
| E. coli *bla* $_{MUN-1}$ | 1 | 4E+06 | 4E+06 | - | fimB | 1-603/603 | =============== | 0/0 | 100.00 | 98.18 | Septicoli | NP_757239 | (fimB) Type 1 fimbriae Regulatory protein fimB [Type 1 fimbriae (VF0221)] [Escherichia coli CFT073]---adhesin | |
| E. coli *bla* $_{MUN-1}$ | 1 | 4E+06 | 4E+06 | - | entD | 1-132/771 | ===...../...... | 02-déc | 15.69 | 84.96 | Septicoli | NP_752599 | (entD) phosphopantetheinyl transferase component of enterobactin synthase multienzyme complex [Enterobactin (VF0228)] [Escherichia coli CFT073]---iron acquisition | |
| E. coli *bla* $_{MUN-1}$ | 1 | 4E+06 | 4E+06 | + | entD | 1-136/771 | ===...../...... | 02-févr | 17.51 | 83.21 | Septicoli | NP_752599 | (entD) phosphopantetheinyl transferase component of enterobactin synthase multienzyme complex [Enterobactin (VF0228)] [Escherichia coli CFT073]---iron acquisition | |
| E. coli *bla* $_{MUN-1}$ | 1 | 4E+06 | 4E+06 | - | entD | 2-114/771 | ===...../...... | 02-févr | 14.53 | 88.60 | Septicoli | NP_752599 | (entD) phosphopantetheinyl transferase component of enterobactin synthase multienzyme complex [Enterobactin (VF0228)] [Escherichia coli CFT073]---iron acquisition | |
| E. coli *bla* $_{MUN-1}$ | 1 | 4E+06 | 4E+06 | + | entD | 6-129/771 | ===...../...... | 02-mars | 15.95 | 81.75 | Septicoli | NP_752599 | (entD) phosphopantetheinyl transferase component of enterobactin synthase multienzyme complex [Enterobactin (VF0228)] [Escherichia coli CFT073]---iron acquisition | |
| E. coli *bla* $_{MUN-1}$ | 1 | 4E+06 | 4E+06 | + | espX5 | 1-1293/1293 | =============== | 0/0 | 100.00 | 97.14 | Septicoli | NP_290699 | (espX5) Type III secretion system effector EspX5 [LEE encoded T3SS (SS020)] [Escherichia coli O157:H7 str. EDL933]---toxin | |
| E. coli *bla* $_{MUN-1}$ | 1 | 4E+06 | 4E+06 | - | espX4 | 126-1581/1581 | .=======/====== | 02-févr | 92.03 | 93.75 | Septicoli | NP_290672 | (espX4) Type III secretion system effector EspX4 [LEE encoded T3SS (SS020)] [Escherichia coli O157:H7 str. EDL933]---toxin | |
| E. coli *bla* $_{MUN-1}$ | 1 | 4E+06 | 4E+06 | - | espX4 | 1-134/1581 | ==............ | 0/0 | 8.48 | 88.06 | Septicoli | NP_290672 | (espX4) Type III secretion system effector EspX4 [LEE encoded T3SS (SS020)] [Escherichia coli O157:H7 str. EDL933]---toxin | |
| E. coli *bla* $_{MUN-1}$ | 1 | 4E+06 | 4E+06 | + | espL4 | 1-2187/2187 | =============== | 0/0 | 100.00 | 97.35 | Septicoli | NP_290644 | (espL4) Type III secretion system effector EspL4 [LEE encoded T3SS (SS020)] [Escherichia coli O157:H7 str. EDL933]---toxin | |
| E. coli *bla* $_{MUN-1}$ | 1 | 5E+06 | 5E+06 | - | entD | 15-124/771 | ===...../...... | 02-févr | 14.14 | 90.09 | Septicoli | NP_752599 | (entD) phosphopantetheinyl transferase component of enterobactin synthase multienzyme complex [Enterobactin (VF0228)] [Escherichia coli CFT073]---iron acquisition | |
| E. coli *bla* $_{MUN-1}$ | 1 | 5E+06 | 5E+06 | - | entD | 1-113/771 | ===...../...... | 01-janv | 14.53 | 88.50 | Septicoli | NP_752599 | (entD) phosphopantetheinyl transferase component of enterobactin synthase multienzyme complex [Enterobactin (VF0228)] [Escherichia coli CFT073]---iron acquisition | |
| E. coli *bla* $_{MUN-1}$ | 1 | 5E+06 | 5E+06 | + | entD | 1-131/771 | ===...../...... | 02-févr | 16.86 | 87.88 | Septicoli | NP_752599 | (entD) phosphopantetheinyl transferase component of enterobactin synthase multienzyme complex [Enterobactin (VF0228)] [Escherichia coli CFT073]---iron acquisition | |
| E. coli *bla* $_{MUN-1}$ | 1 | 5E+06 | 5E+06 | - | entD | 1-137/771 | ===...../...... | 02-févr | 17.64 | 84.78 | Septicoli | NP_752599 | (entD) phosphopantetheinyl transferase component of enterobactin synthase multienzyme complex [Enterobactin (VF0228)] [Escherichia coli CFT073]---iron acquisition | |
| E. coli *bla* $_{MUN-1}$ | 1 | 5E+06 | 5E+06 | + | aslA | 1-1428/1428 | ========/====== | 01-janv | 100.00 | 93.56 | Septicoli | AAG10151 | (aslA) putative arylsulfatase [AslA (VF0238)] [Escherichia coli O18:K1:H7 str. RS218]---invasin | |
| E. coli *bla* $_{MUN-1}$ | 3 | 46600 | 47331 | + | traT | 1-732/732 | =============== | 0/0 | 100.00 | 98.09 | Septicoli | J01769.1:403-1134 | Plasmid R100 (from E.coli) complement resistance (traT) gene---protectin | |
