## Supplementary Table 4 for "Inter-phylum circulation of a beta-lactamase - encoding gene: a rare but observable event"

Supplementary Table 4: Hits obtained analyzing the blaMUN-1 gene distribution using BLASTN against RefSeq genomes database.

| Species | RefSeq Genomes entries | % identity | Query cover | Accession numbers | max copies /AN |
|---|---|---|---|---|---|
| *Alistipes shahii* | 31 | 100 | 100 | NZ_JADMNV010000001.1 | 1 |
| *Alistipes putredinis* | 75 | 100 | 100 | NZ_CAUFBO010000004.1 | 1 |
| *Bacteroides caccae* | 99 | 100 | 100 | NZ_RCXH01000036.1; NZ_VVYI01000057.1; NZ_VVYJ01000032.1 | 1 |
| *Bacteroides eggerthii* | 54 | 100 | 100 | NZ_RCXL01000057.1; NZ_VVZY01000116.1; NZ_VVZX01000033.1 | 1 |
| *Bacteroides fragilis* | 566 | 100 | 100 | NZ_VOHY01000006.1; NZ_JANGCS010000017.1; NZ_JAJCKU010000030.1 | 1 |
| *Bacteroides ovatus* | 288 | 100 | 100 | NZ_JAHYNG010000035.1; NZ_JAQEVX010000058.1 | 1 |
| *Bacteroides salyersiae* | 58 | 99.88-100 | 100 | NZ_WCKY01000001.1; NZ_WCKW01000090.1; NZ_WCKX01000075.1; NZ_JABFHZ010000002.1; NZ_CP081902.1 | 1 |
| *Bacteroides stercoris* | 98 | 100 | 100 | NZ_CABOGI010000053.1; NZ_JAQNVU010000011.1; NZ_QSSP01000053.1; NZ_QROW01000004.1; NZ_JAQNVT010000010.1 | 1 |
| *Bacteroides thetaiotaomicron* | 328 | 100 | 100 | NZ_JAHYLL010000075.1; NZ_JAHYPH010000024.1; NZ_JADPBE010000005.1; NZ_JAQNVD010000005.1; NZ_JANUPG010000001.1 | 1 |
| *Bacteroides uniformis* | 365 | 100 | 100 | NZ_WCUR01000119.1; NZ_WCUP01000009.1; NZ_WCUQ01000001.1; NZ_JANUJA010000001.1; NZ_AP019725.1; NZ_AP019724.1 | 6 |
| *Bacteroides sp.* | 53 | 100 | 100 | NZ_JAPQKZ010000100.1; NZ_CP083673.1; NZ_CAAFEE010000290.1 | 1 |
| *Bacteroides xylanisolvens* | 195 | 99.88-100 | 100 | NZ_WDEE01000003.1; NZ_WDEF01000003.1; NZ_JAHOJA010000006.1 | 1 |
| *Barnesiella propionica* | 1 | 71.93 | 96 | NZ_JAOQJK010000002.1 | 1 |
| *Butyricimonas faecihominis* | 8 | 100 | 100 | NZ_JAQEXV010000004.1 | 1 |
| *Odoribacter splanchnicus* | 52 | 100 | 82-100 | NZ_JADNDE010000093.1; NZ_JADNIQ010000002.1; NZ_QSCO01000022.1 | 1 |
| *Parabacteroides distasonis* | 253 | 99.88-100 | 100 | NZ_JAHONS010000002.1; NZ_JAHONT010000002.1; NZ_JAHYMP010000020.1; NZ_JAJCJX010000007.1; NZ_JAJCNE010000002.1; NZ_WKMM01000035.1; NZ_JAQEXU010000002.1; NZ_JAQMQB010000055.1; NZ_JAQMQC010000053.1; NZ_AP019729.1; NZ_BQOD01000001.1; NZ_BQOC01000001.1; NZ_JAHYMB010000033.1; NZ_CP103256.1 | 1 |
| *Parabacteroides faecis* | 5 | 100 | 100 | NZ_JACRTM010000033.1 | 1 |
| *Parabacteroides goldsteinii* | 102 | 100 | 100 | NZ_BQND01000001.1; NZ_BQNC01000001.1 | 2 |
| *Parabacteroides johnsonii* | 36 | 100 | 100 | NZ_JAASIA010000002.1; NZ_JANUNE010000002.1; NZ_JANUNE010000003.1 | 3 |
| *Parabacteroides merdae* | 126 | 100 | 100 | NZ_JADNHS010000020.1; NZ_JADMOA010000018.1; NZ_JAQDNL010000017.1; NZ_JAQMOI010000016.1; NZ_JAQMOJ010000018.1 | 1 |
| *Parabacteroides sp.* | | | | NZ_KQ236102.1; NZ_JAMOKM010000041.1; NZ_JAOEGA010000001.1; NZ_QTMV01000009.1 | 1 |
| *Leyella stercorea* | 49 | 99.88-100 | 100 | NZ_JAIJUX010000025.1; NZ_QRNO01000005.1; NZ_CABJDY010000005.1 | 1 |
| *Paraprevotella clara* | 27 | 100 | 100 | NZ_AP025941.1 | 2 |
| *Prevotellamassilia timonensis* | 3 | 100 | 100 | NZ_LT629842.1; NZ_LT629840.1; NZ_LT629830.1; NZ_LT629845.1 | 1 |
| *Phocaeicola massiliensis* | 27 | 100 | 100 | NZ_JADMTU010000014.1; NZ_JAQCSN010000016.1; NZ_JAQCTA010000014.1 | 1 |
| *Phocaeicola dorei* | 114 | 100 | 100 | NZ_BQOB01000001.1; NZ_BQOA01000001.1; NZ_JAKNHN010000004.1; NZ_JADMOY010000018.1; NZ_JADNHW010000026.1; NZ_JADMSQ010000098.1; NZ_WQQY01000001.1; NZ_JAKNHU010000046.1; NZ_JAJCJS010000047.1; NZ_JAHYPM010000077.1; NZ_JAQDJF010000007.1; NZ_JAQDJE010000006.1; NZ_JAFBJH010000021.1; NZ_JAFBJF010000021.1 | 1 |
| *Phoceicola vulgatus* | 544 | 96.98-100 | 100 | NZ_QSSN01000021.1; NZ_QSBO01000007.1; NZ_JADPEZ010000016.1; NZ_QRMN01000001.1; NZ_QROU01000027.1; NZ_JAKKXJ010000056.1; NZ_JAKKXN010000001.1; NZ_JAKKXS010000056.1; NZ_JAKKWW010000002.1; NZ_JAKKXI010000074.1; NZ_JAKKXB010000001.1; NZ_JANUTN010000007.1; NZ_JAHYRE010000075.1; NZ_QRYD01000047.1; NZ_QRYF01000060.1; NZ_JAHOIR010000010.1; NZ_JAHOIT010000010.1; NZ_JAHOIU010000008.1; NZ_JAHOIM010000010.1; NZ_JAHOIW010000010.1; NZ_JAHOIX010000010.1; NZ_JAHOIV010000010.1; NZ_JAHOIS010000010.1; NZ_JAHOIL010000009.1; NZ_JAHOIN010000010.1 | 2 |
| *Porphyromonas somerae* | 4 | 100 | 100 | NZ_AP025559.1 | 1 |
| *Sutterella wadsworthensis* | 62 | 100 | 100 | NZ_JAANXS010000022.1 | 1 |
