## Supplementary Table 5 for "Inter-phylum circulation of a beta-lactamase - encoding gene: a rare but observable event"

Supplementary Table 5: Hits obtained analyzing the blaMUN-1 gene distribution using MGnify.

| Accession | Catalogue | Type | Taxonomy | K-mers in query | K-mers found in genome | % K-mers found |
|---|---|---|---|---|---|---|
| MGYG000003681 | human-gut-v2-0-1 | Isolate | Bacteroides stercoris | 798 | 798 | 100 |
| MGYG000303511 | pig-gut-v1-0 | MAG | Onthomorpha sp016296345 | 798 | 798 | 100 |
| MGYG000000215 | human-gut-v2-0-1 | Isolate | Prevotella stercorea | 798 | 775 | 97.12 |
| MGYG000003252 | human-gut-v2-0-1 | MAG | Bacteroides sp900761785 | 798 | 775 | 97.12 |
