## Supplementary Table 6 for "Inter-phylum circulation of a beta-lactamase - encoding gene: a rare but observable event"

Supplementary Table 6: Hits obtained analyzing the MUN-1 protein distribution using the GMGC catalog.

| Unigene | E-value | Complete | Habitat | Taxon (predicted) |
|---|---|---|---|---|
| GMGC10.047_051_980.UNKNOWN | 4.76e-154 | 1 | human gut,human nose,human oral,human skin,human vagina,mouse gut | Prevotellamassilia timonensis (species) |
| GMGC10.047_909_394.UNKNOWN | 4.93e-143 | 1 | human gut | Prevotellamassilia timonensis (species) |
| GMGC10.051_471_042.UNKNOWN | 4.93e-143 | 1 | human gut | Prevotellamassilia timonensis (species) |
| GMGC10.302_260_782.UNKNOWN | 8.71e-140 | 1 | human gut | Prevotellamassilia timonensis (species) |
| GMGC10.203_264_893.UNKNOWN | 6.9e-137 | 1 | human gut | Prevotellamassilia timonensis (species) |
| GMGC10.207_492_799.UNKNOWN | 4.49e-120 | 1 | human gut | Prevotellamassilia timonensis (species) |
| GMGC10.306_704_321.UNKNOWN | 3.45e-112 | 1 | human gut | Prevotellamassilia timonensis (species) |
| GMGC10.052_355_696.UNKNOWN | 1.31e-103 | 1 | cat gut,human gut | Prevotellamassilia timonensis (species) |
| GMGC10.190_096_469.UNKNOWN | 5.16e-100 | 1 | human gut,pig gut | Prevotellamassilia timonensis (species) |
| GMGC10.209_122_821.UNKNOWN | 2.56e-99 | 1 | human gut | Prevotellamassilia timonensis (species) |
| GMGC10.206_835_329.UNKNOWN | 3.13e-97 | 1 | human gut | Parabacteroides johnsonii DSM 18315 (species) |
| GMGC10.050_469_855.UNKNOWN | 7.72e-96 | 1 | human gut | Prevotellamassilia timonensis (species) |
| GMGC10.208_088_023.UNKNOWN | 1.51e-91 | 1 | human gut | Parabacteroides johnsonii DSM 18315 (species) |
| GMGC10.055_103_744.UNKNOWN | 1.32e-87 | 1 | human gut | Parasutterella excrementihominis YIT 11859 (species) |
| GMGC10.206_640_283.UNKNOWN | 6.55e-87 | 1 | human gut | Alistipes sp. CAG:831 (species) |
| GMGC10.001_916_490.UNKNOWN | 1.23e-85 | 0 | dog gut | Parabacteroides goldsteinii (species) |
| GMGC10.050_503_506.UNKNOWN | 2.58e-83 | 1 | human gut | Bacteroides sp. CAG:144 (species) |
| GMGC10.000_544_401.UNKNOWN | 1.67e-82 | 1 | dog gut,human gut | Parasutterella excrementihominis YIT 11859 (species) |
| GMGC10.147_033_135.UNKNOWN | 1.85e-81 | 1 | human gut,mouse gut | Parasutterella excrementihominis YIT 11859 (species) |
| GMGC10.056_912_393.UNKNOWN | 2.41e-81 | 1 | human gut,mouse gut | Parasutterella excrementihominis YIT 11859 (species) |
| GMGC10.210_339_896.UNKNOWN | 5.37e-81 | 1 | human gut,mouse gut | Parasutterella excrementihominis YIT 11859 (species) |
| GMGC10.054_684_518.UNKNOWN | 9.16e-81 | 1 | human gut | Bacteroides sp. CAG:144 (species) |
| GMGC10.055_669_095.UNKNOWN | 1.2e-80 | 1 | human gut,mouse gut | Parasutterella excrementihominis YIT 11859 (species) |
| GMGC10.306_210_464.UNKNOWN | 1.01e-79 | 1 | human gut | Parasutterella excrementihominis YIT 11859 (species) |
| GMGC10.310_770_158.UNKNOWN | 9.5e-70 | 1 | cat gut,human gut,human nose,human vagina | Bacteroides sp. CAG:20 (species) |
| GMGC10.306_122_048.UNKNOWN | 5.21e-68 | 1 | human gut | Bacteroides timonensis (species) |
| GMGC10.184_778_431.UNKNOWN | 1.52e-67 | 1 | human gut,human oral | Bacteroides timonensis (species) |
| GMGC10.309_096_849.UNKNOWN | 1.28e-66 | 1 | human gut,human oral,mouse gut | Bacteroides intestinalis (species) |
| GMGC10.207_343_478.UNKNOWN | 1.68e-66 | 0 | human gut,human oral,mouse gut | Bacteroides timonensis (species) |
| GMGC10.207_362_373.UNKNOWN | 1.68e-66 | 0 | human gut,human oral,mouse gut | Bacteroides timonensis (species) |
| GMGC10.055_127_397.UNKNOWN | 2.86e-66 | 1 | human gut | Bacteroides stercorirosoris (species) |
| GMGC10.032_188_571.UNKNOWN | 1.09e-65 | 1 | - | Bacteroides stercorirosoris (species) |
| GMGC10.177_802_870.UNKNOWN | 1.62e-61 | 1 | mouse gut | Bacteroidales (order) |
| GMGC10.298_506_683.UNKNOWN | 2.12e-61 | 1 | human gut | Odoribacter laneus CAG:561 (species) |
| GMGC10.207_032_025.UNKNOWN | 2.77e-61 | 1 | human gut | Parabacteroides timonensis (species) |
| GMGC10.297_552_894.UNKNOWN | 4.73e-61 | 0 | human gut | Bacteroidales (order) |
| GMGC10.207_524_736.UNKNOWN | 8.06e-61 | 1 | human gut | Parabacteroides timonensis (species) |
| GMGC10.057_236_757.UNKNOWN | 2.35e-60 | 1 | human gut | Parabacteroides timonensis (species) |
| GMGC10.050_713_307.UNKNOWN | 1.52e-59 | 1 | human gut | Bacteroides sp. Marseille-P3108 (species) |
| GMGC10.174_742_840.UNKNOWN | 7.54e-59 | 1 | human gut,mouse gut | Bacteroides (genus) |
| GMGC10.184_003_774.UNKNOWN | 7.54e-59 | 1 | human gut,mouse gut | Bacteroides congonensis (species) |
| GMGC10.306_142_568.UNKNOWN | 1.68e-58 | 1 | cat gut,dog gut,human gut,human nose,human oral,human skin,human vagina,mouse gut | Bacteroides vulgatus (species) |
| GMGC10.256_862_127.SP_0010 | 2.2e-58 | 1 | human gut | Bacteria (superkingdom) |
| GMGC10.206_992_817.UNKNOWN | 2.87e-58 | 1 | human gut,mouse gut | Bacteroides congonensis (species) |
| GMGC10.288_000_481.UNKNOWN | 1.09e-57 | 0 | cat gut,dog gut,human gut,human nose,human oral,human skin,human vagina,mouse gut | Bacteroides dorei 5_1_36/D4 (species) |
| GMGC10.177_589_833.UNKNOWN | 1.86e-57 | 1 | mouse gut | Bacteroides (genus) |
| GMGC10.211_900_450.UNKNOWN | 4.14e-57 | 1 | human gut | Parabacteroides timonensis (species) |
| GMGC10.050_299_762.UNKNOWN | 5.41e-57 | 1 | human gut,mouse gut | Bacteroides acidifaciens JCM 10556 (species) |
| GMGC10.178_223_713.UNKNOWN | 5.41e-57 | 1 | human gut,mouse gut | Bacteroides acidifaciens JCM 10556 (species) |
| GMGC10.177_403_137.UNKNOWN | 7.06e-57 | 1 | mouse gut | Alistipes obesi (species) |
