## Supplementary Table 1 for "Inter-phylum circulation of a beta-lactamase - encoding gene: a rare but observable event"

Supplementary Table 1: Differences in terms of contigs obtained after assembly using Unicycler between white and grey colonies after molecular characterization.

| | Chromosome | | incFII | | p0111 | |
|---|---|---|---|---|---|---|
| | size | *n* *bla* $_{MUN-1}$ | size (bp) | *n* *bla* $_{MUN-1}$ | size (bp) | *n* *bla* $_{MUN-1}$ |
| White | 4,762,657 | 2 | 60,123 | 0 | 127,245 | 2 |
| Grey | 4,764,212 | 2 | 60,123 | 0 | - | - |
