## Supplementary Table 2 for "Inter-phylum circulation of a beta-lactamase - encoding gene: a rare but observable event"

Supplementary Table 2: Antibiotic resistance genes (ARGs) found in the chromosome of the E. coli strain using ResFinder (v4.0).

| ARG Family | ARG name | Description | Resistance to | Contig number | Identity | Coverage | Access number |
|---|---|---|---|---|---|---|---|
| ANT | *aadA5* | ANT(3'')-Ia family aminoglycoside nucleotidyltransferase AadA5 | Aminoglycosides | bacteroides_hybrid_03923 | 100.0 | 99.6 | NG_047357.1.1 |
| APH | *aph(3'')-Ib* | aminoglycoside O-phosphotransferase APH(3'')-Ib | Aminoglycosides | bacteroides_hybrid_03940 | 100.0 | 99.6 | NG_047413.1.1 |
| APH | *aph(6)-Id* | aminoglycoside O-phosphotransferase APH(6)-Id | Aminoglycosides | bacteroides_hybrid_03941 | 100.0 | 99.6 | NG_047464.1.1 |
| APH | *aph(3')-Ia* | aminoglycoside O-phosphotransferase APH(3')-Ia | Aminoglycosides | bacteroides_hybrid_03943 | 100.0 | 99.6 | NG_047430.1.1 |
| Cat | *catA1* | type A-1 chloramphenicol O-acetyltransferase | Chloramphenicol | bacteroides_hybrid_03916 | 99.3 | 65.9 | NG_051704.1.1 |
| Class A beta-lactamase | $bla_{\text{TEM-1}}$ | class A broad-spectrum beta-lactamase TEM-1 | Beta-lactams | bacteroides_hybrid_03934 | 100.0 | 99.7 | NG_050145.1.1 |
| Class C beta-lactamase | $bla_{\text{EC}}$ | BlaEC family class C beta-lactamase | Beta-lactams | bacteroides_hybrid_00194 | 100.0 | 99.7 | NG_047494.1.1 |
| Dfr | *dfrA17* | trimethoprim-resistant dihydrofolate reductase DfrA17 | Trimethoprim | bacteroides_hybrid_03922 | 99.4 | 99.4 | NG_047709.1.1 |
| Mph | *mph(A)* | Mph(A) family macrolide 2'-phosphotransferase | Macrolides | bacteroides_hybrid_03932 | 100.0 | 99.7 | NG_047985.1.1 |
| Sul | *sul1* | sulfonamide-resistant dihydropteroate synthase Sul1 | Sulphonamides | bacteroides_hybrid_03925 | 99.6 | 99.6 | NG_048081.1.1 |
| Sul | *sul2* | Sulfonamide-resistant dihydropteroate synthase Sul2 | Sulphonamides | bacteroides_hybrid_03939 | 100.0 | 99.6 | NG_048118.1.1 |
| Tet efflux | *tet(B)* | tetracycline efflux MFS transporter Tet(B) | Tetracyclines | bacteroides_hybrid_03513 | 100.0 | 99.8 | NG_048163.1.1 |
